# Systemic Multi-Omics Analysis Reveals Interferon Response Heterogeneity and Links Lipid Metabolism to Immune Alterations in Severe COVID-19

**DOI:** 10.1101/2025.03.14.643374

**Authors:** Ronaldo Lira, Anoop T Ambikan, Axel Cederholm, Sefanit Rezene, Flora Mikaeloff, Sara Svensson Akusjärvi, Ahmet Yalcinkaya, Xi Chen, Maike Sperk, Maribel Aranda-Guillén, Hampus Nordqvist, Carl Johan Treutiger, Nils Landegren, Ujjwal Neogi, Soham Gupta

## Abstract

The immune response to SARS-CoV-2 infection is highly heterogeneous, and interferon (IFN)-stimulated genes (ISGs) play a central but context-dependent role in antiviral defense and immune dysregulation. To investigate how ISG heterogeneity relates to immune and metabolic states, we performed an integrated analysis of whole-blood transcriptomics, plasma proteomics, metabolomics, and immune activation markers in hospitalized COVID-19 patients and COVID-negative healthy controls and covalescent individuals. Patients segregated into low (LIS), moderate (MIS), and high (HIS) ISG expression endotypes, largely independent of clinical severity. While high ISG expression was associated with systemic inflammation and innate immune activation, severe disease within the HIS endotype was characterized by marked metabolic perturbations, including depletion of tricarboxylic acid cycle intermediates and multiple lipid classes involved in membrane integrity and immunometabolic signaling. Plasma-transfer assays demonstrated that plasma from severe HIS patients impaired neutrophil and monocyte activation ex vivo, indicating functional attenuation of innate immune responses despite elevated ISG expression. These metabolic alterations correlated with reduced immune activation, supporting the existence of an interferon-associated immune-metabolic axis that constrains immune functionality in severe disease. Although type I IFN neutralization was detected in a subset of patients with IFN antigen reactivity, these samples did not account for ISG heterogeneity or disease severity. Together, these findings show that high ISG expression defines a transcriptional endotype permissive for inflammation but insufficient for effective immune function, highlighting the importance of immune–metabolic context in shaping COVID-19 disease outcomes.

## Introduction

The coronavirus disease 2019 (COVID-19) is caused by the severe acute respiratory syndrome coronavirus 2 (SARS-CoV-2) which exhibits a wide range of clinical manifestations, from asymptomatic infection to severe respiratory distress with systemic complications [1]. A key feature of severe infection is hyperinflammation driven by dysregulated immune signaling caused by proinflammatory cytokines, including alterations in interferon (IFN) responses [2]. Interferons (IFN) are specialized type of cytokines that mediate antiviral defense upon viral RNA recognition by receptors such as RIG-I or MDA-5 receptors [3]. IFNs induce the interferon-stimulated genes (ISGs) through a signaling cascade, activating three major IFN-classes: type I IFN (IFN-α, -β, -ε, -κ, -ω), type II (IFN-γ), and type III (IFN-λ) [4]. While these IFNs signal through distinct receptors, they share downstream ISG activation, with significant overlap between type I and type III IFN responses [5, 6].

Interferons (IFNs) are among the earliest antiviral responses, with an early and robust response being critical for effective viral clearance and protection against SARS-CoV-2 infection [7]. However, genetic defects and presence of specific IFN-blocking autoantibodies can dampen their antiviral role and cause an inability to control SARS-CoV-2 replication, leading to a more severe disease [8, 9]. While some studies report a suppressed IFN response in severe cases [10], others describe an elevated ISG expression with heightened inflammation, which correlates with worse disease outcomes [11]. SARS-CoV-2 is known to actively disrupt innate immune responses, contributing to dysregulated IFN signaling [12]. In murine models of human coronavirus infection, a delayed type I IFN response has been linked to excessive monocyte and macrophage activation, driving proinflammatory cytokine storms [4]. Beyond the immune response, metabolic alterations have emerged as critical determinants of COVID-19 severity. IFN signaling can influence lipid metabolism, glycolysis, and amino acid pathways [13]. These metabolic shifts can, in turn, modulate immune cell function and inflammatory outcomes [13]. Furthermore, the dysregulation in IFN response can induce both autoimmune and non-autoimmune inflammatory responses [14]. Autoantibodies against type I IFNs have been implicated in modifying IFN signaling in a subset of patients, although their precise role in disease progression remains complex and context-dependent [9, 15]. Thus, the efficient viral clearance and disease outcome hangs in a balance between IFN response and inflammation. There is considerable multifariousness in the IFN response and understanding both the protective and aberrant responses can aid in the development of immunomodulatory therapeutic strategies.

Beyond acute infection, IFN response heterogeneity extends beyond COVID-19 severity, influencing post-viral immune dysregulation, persistent inflammation, and metabolic dysfunction. Emerging evidence suggests that dysregulated ISG expression may contribute to post-viral syndromes such as long COVID, where prolonged immune activation and metabolic shifts are implicated in sustained fatigue, neurological symptoms, and cardiovascular complications [16]. Additionally, IFN-driven metabolic reprogramming, particularly disruptions in lipid and amino acid metabolism, may underlie broader immune dysfunctions, including autoimmune-like sequelae [17]. Understanding the interplay between IFN signaling, immune metabolism, and inflammatory resolution is therefore critical not only for COVID-19 but also for broader applications in chronic inflammatory and post-viral syndromes.

In this study, we investigate biological perturbations associated with ISG expression patterns in COVID-19 patients, stratifying individuals based on ISG-driven clustering (low-ISG score: LIS, moderate-ISG score: MIS, and high-ISG score: HIS). Using multi-omics data (plasma proteomics and metabolomics), IFN antigen-reactivity screening, and immune activation analyses, we explore how IFN response heterogeneity is linked to immune and metabolic alterations. Our findings reveal a stochastic relationship between ISG expression and immune cell activation, along with significant perturbations in lipid and amino acid metabolism. These insights offer a framework for understanding the complex interplay between IFN signaling, immune responses, and metabolic adaptation in COVID-19, which could aid in personalized treatment strategies.

## Results

### IFN-signatures in COVID-19 disease severity

Interferons (IFNs) induce interferon-stimulated genes (ISGs) with antiviral functions, and dysregulated IFN responses have been linked to COVID-19 severity. Although SARS-CoV-2 primarily infects respiratory tissues, whole-blood profiling captures systemic interferon signaling and circulating immune-metabolic states that may modulate disease course. To investigate the link between IFN signaling and disease severity, we analysed whole-blood RNA-seq data from 21 COVID-negative healthy controls (HC), 10 Convalescent individuals (Conv), 26 Mild and 11 Severe COVID-19 patients (Figure 1A) previously described by us [18]. Transcriptomic analyses focused on genes involved in the type I IFN (IFN-α/β) signaling pathway (R-HSA-909733), type II IFN (IFN-γ) signaling pathway (R-HSA-877300), and antiviral ISG pathway (R-HSA-1169410) (Table S1), with overlapping ISGs across pathways. Both the mild and severe patient groups showed a significant increase in ISG expression, predominantly within the type I IFN signaling (FDR<0.05, Figure 1B). To capture patient-specific IFN responses, we computed ISG scores for type I, type II, and antiviral signaling pathways by summing gene-wise Z-scores within a given pathway [19] (Table S2). A significant increase in ISG score for all three pathways was observed in mild disease compared to healthy, while no difference was seen between the mild and severe groups (Figure 1C, Mann Whitney U test, *p*-value<0.05). Despite similar aggregate ISG scores, visualization of ISG expression revealed distinct gene expression patterns between mild and severe cases, indicating underlying heterogeneity (Figure 1B). All the severe cases in our cohort were male and no sex-specific separation of ISG-scores was observed within the mild group.

**Figure 1:**
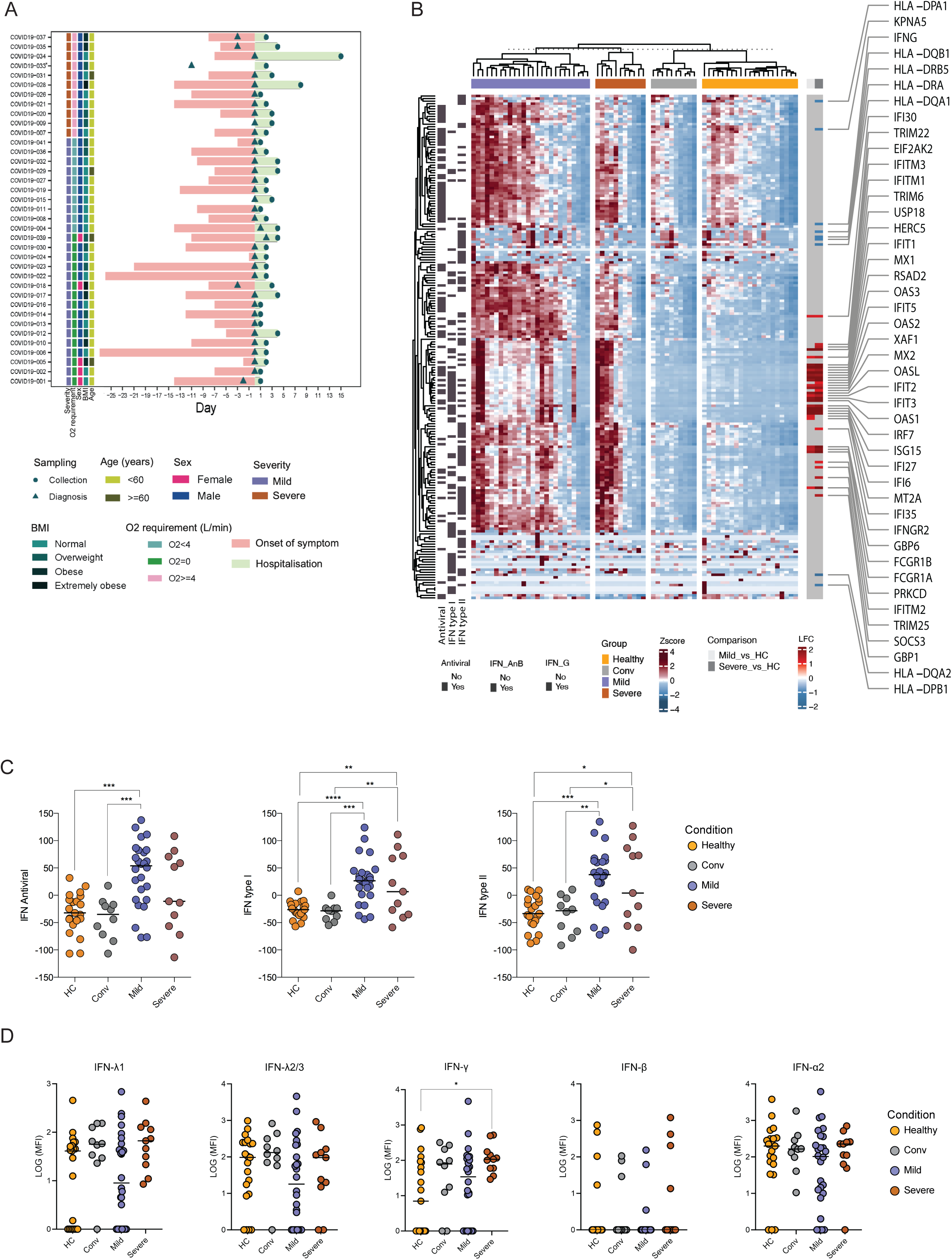
Transcriptomic and plasma IFN analysis of COVID-negative healthy controls (HC; n = 21), convalescent individuals (Conv; n = 10), mild COVID-19 patients (n = 26), and severe COVID-19 patients (n = 11). **A)** Schematic figure showing the clinical characteristics of COVID-19 infected individuals. **B)** Heatmap of the interferon (IFN) stimulated genes (ISG) profile of the clinical cohort from transcriptomic data. **C)** Dot plots of the ISG score for type I ISGs, type II ISGs, and antiviral ISGs in the clinical groups separately. **D)** Plasma levels of IFN-λ1, IFN-λ2/3, IFN-γ, IFN-β, and IFN-α2 measured by LEGENDplex assay expressed as median fluorescent intensity (MFI). Statistical significance was measured using Kruskal-Wallis test with significance * *p*<0.05, ** *p*<0.01, *** *p*<0.001, and **** *p*<0.0001.

IFNs activate ISGs via the JAK-STAT pathway. To assess whether ISG expression reflected circulating IFN levels, we examined IFN gene expression and plasma IFN concentrations. At the transcript level, most IFN gene were expressed at low or undetectable levels (Transcripts per million, TPM <1), except for IFN-λ1 and IFN-γ, showing reduced expression in severe cases (Figure S1). Plasma IFN-I (IFN-α2 and IFN-β), IFN-II (IFN-γ), and IFN-III (IFN-λ1 and IFN-λ2/3) were measured using a FACS-based LEGENDplex assay. Due to low concentrations, median fluorescence intensity (MFI) was used for analysis. No significant differences in most circulating IFN levels were observed across patient groups (Kruskal-Wallis test, FDR > 0.05, Figure 1D, Table S3). However, IFN-γ was higher in severe COVID-19 compared with HC (adjusted p = 0.0427).

### COVID-19 patient stratification based on the ISG expression

ISG expression showed marked heterogeneity between clinical groups. While ISG scores were uniformly distributed among mild cases, severe cases displayed a bimodal distribution, with five patients exhibiting high ISG scores and six showing near-baseline levels (Figure 1C). This pattern prompted re-categorization of all samples based on their transcriptomic ISG profiles. To address this heterogeneity, we performed a robust hierarchical consensus clustering of the patients based on ISGs expression profile. This approach segregated subjects into three clusters, supported by a consensus matrix and visualized by PCA analysis (Figures 2A, Figure S2). Clusters were defined based on ISG scores across the three IFN pathways and designated as low-ISG score (LIS), moderate-ISG score (MIS), and high-ISG score (HIS), with LIS resembling the healthy control profile (Mann Whitney U test, *p*-value<0.05, Figure 2B-2C). The alignment of these clusters with previous clinical categorizations is displayed in Figure 2B. Notably, ISG-defined clustering did not fully align with clinical severity. While most mild cases clustered within MIS and HIS, severe cases were distributed across both LIS and HIS (Figure 2B), indicating that severe COVID-19 can occur in the context of either low or high systemic ISG expression. This dissociation suggests that ISG magnitude alone is insufficient to explain disease severity and points to the existence of distinct immunological endotypes among hospitalized patients. To explore the biological basis of this heterogeneity, we next examined whether hospitalized patients with severe disease differed depending on their ISG-defined endotype, focusing on systemic inflammation, immune activation, and metabolic perturbations.

**Figure 2:**
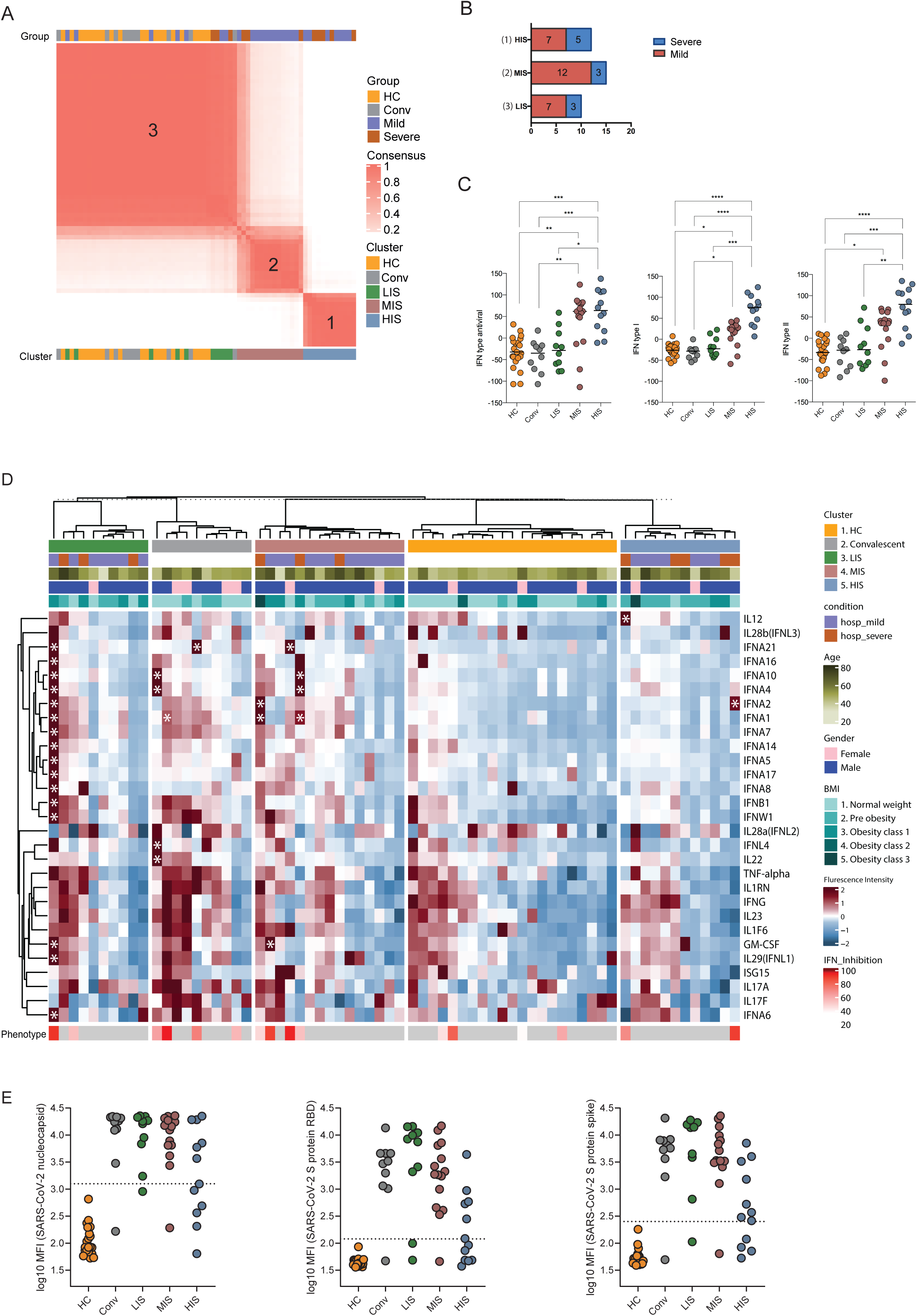
Clustering of COVID-19 patients based on ISG transcriptomic profiles and detection of IFN antigen reactivity. **A)** Heatmap of the consensus clustering matrix depicting patient clustering based on ISG expression. Patients are displayed along both rows and columns, with clinical conditions (top) and ISG-based clusters (bottom) indicated. Consensus values range from 0 (no co-clustering) to 1 (consistent co-clustering) and are color-coded from white to red. Clinical severity (mild/severe) and ISG-defined clusters (LIS/MIS/HIS) represent independent classifications; color coding reflects severity distribution within each cluster. **B)** Bar plot showing the distribution of clinically categorized subjects within each ISG-defined cluster (low-ISG: LIS, moderate-ISG: MIS, and high-ISG: HIS). **C)** Box plots illustrating the antiviral, type I IFN, and type II IFN ISG scores across different ISG-defined clusters, highlighting differential ISG expression patterns. **D)** Heatmap showing reactivity against recombinant IFN antigens across patient groups and ISG-defined clusters. For each sample–antigen pair, antigen reactivity was assessed by mean fluorescence intensity (MFI), calculated from multiple beads coupled to the same antigen. MFI values were scaled per antigen for visualization. Antigen-reactive samples exceeding the antigen-specific positivity threshold, defined as the mean MFI of healthy controls + 7 SD, are marked with asterisks. The bottom annotation row displays type I IFN neutralization assay results for the corresponding samples where performed. Clinical metadata, including patient condition, ISG-defined cluster, sex, age, and BMI category, are displayed above the heatmap. **E)** Dot plots showing antibody reactivity against SARS-CoV-2 nucleocapsid, spike receptor-binding domain (RBD), and spike antigens across ISG-defined clusters. Each point represents one sample, and antibody reactivity is shown as log10-transformed mean fluorescence intensity (log10 MFI). The horizontal dashed line indicates the antigen-specific positivity threshold, defined as the mean MFI of healthy controls + 7 SD.

### IFN antigen reactivity and neutralizing autoantibodies across ISG-defined patient clusters

Inborn errors of IFN immunity and neutralizing autoantibodies against IFNs have been associated with immune dysregulation and severe COVID-19 [8, 9]. To assess whether IFN antigen reactivity contributed to ISG-defined heterogeneity, we screened patient plasma using an autoantigen array featuring recombinant type I, type II, and type III IFN antigens. Samples exceeding a stringent positivity threshold (mean MFI of healthy controls +7 SD) were considered reactive (marked with asterisks in Figure 2D). Samples with IFN antigen reactivity were detected in a small subset of COVID-19 patients across LIS, MIS, and HIS, as well as in a minority of convalescent samples. Most IFN antigen-reactive COVID-19 patients clustered within MIS, with single cases observed in LIS and HIS. Additional reactivities against GM-CSF and IL-12 were also detected (Figure 2D).

To evaluate the functional impact of IFN-α2 antigen reactivity, neutralization assays were performed on IFN antigen-reactive samples using a dual-luciferase ISRE reporter system. Four of six tested antigen-reactive samples showed >80% neutralization capacity, spanning LIS, MIS, and HIS groups (bottom row, Figure 2D). Notably, only one neutralizing sample corresponded to a severe HIS case, and one convalescent sample also displayed IFN-neutralizing activity. Importantly, no consistent association was observed between IFN antigen reactivity, ISG-defined clustering, or clinical severity. We additionally assessed antibody reactivity against SARS-CoV-2 antigens. HIS patients showed reduced spike and RBD antibody reactivity compared to LIS and MIS, which more closely resembled convalescent profiles (Figure 2E). In contrast, nucleocapsid-directed antibody responses were preserved across ISG groups, suggesting its potential to be an effective immunogen with implications for both vaccine design and diagnostics.

Although a subset of patients showed IFN-neutralizing activity, this did not correlate with ISG-defined clustering or disease severity in this cohort. These findings indicate that IFN antigen reactivity does not account for the ISG-associated immune-metabolic endotypes identified here, but may represent an additional layer of immune dysregulation in a subset of individuals.

### Plasma Proteomics Reveals a Progressive Increase in Inflammation with Higher ISG Expression

To investigate the relationship between ISG expression and systemic inflammation, we analyzed the plasma inflammatory cytokines measured using Olink onco-immunology panel [20] across ISG-defined clusters. Principal Component Analysis (PCA) revealed a clear segregation of samples that mirrored the ISG score-based clustering (Figure 3A). Healthy controls and convalescents clustered closely, while LIS samples did not form a tight cluster with healthy controls or convalescent individuals, but instead displayed a dispersed distribution spanning the space between HC/Conv and MIS COVID-19 patients. Whereas MIS and HIS samples progressively diverged along PC1 (47.04% variance), indicating an increasing systemic inflammatory burden with higher ISG expression.

**Figure 3:**
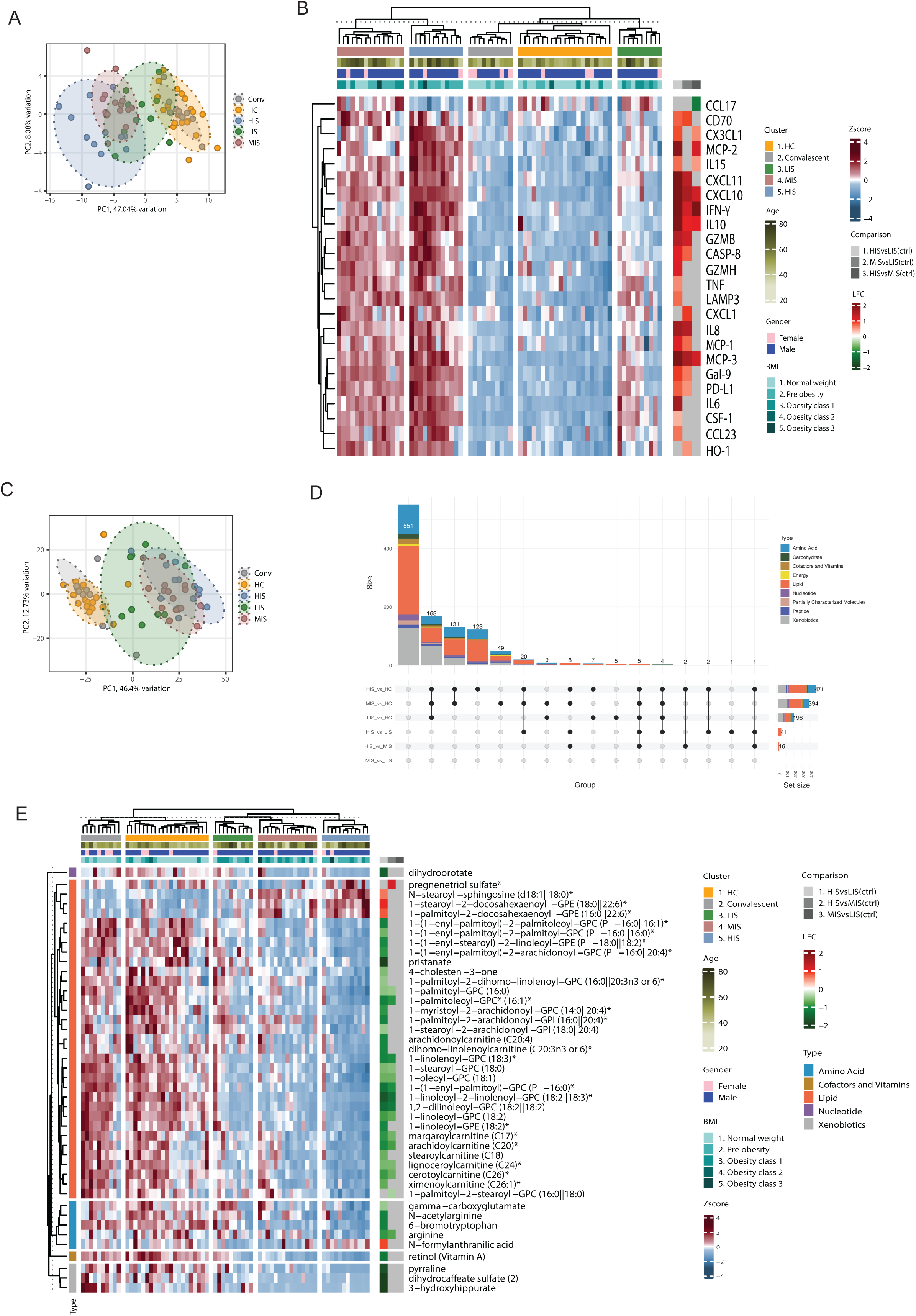
Multi-omics analysis of inflammatory and metabolic perturbations across ISG-defined clusters. Plasma proteomics and metabolomics data were derived from the previously published dataset in reference [19]. **A)** Principal Component Analysis (PCA) plot of Olink proteomic data from healthy controls (HC), convalescent individuals (Conv), and COVID-19 patients stratified into LIS, MIS, and HIS clusters. The first two principal components are displayed, with variance explained indicated on the axes and color-coded cluster categories. **B)** Heatmap of significantly differentially expressed plasma proteins across ISG-defined clusters (LIS, MIS, HIS). The main heatmap displays z-score–normalized protein abundance across individual samples, with hierarchical clustering of patients. Columns represent individual samples, rows represent proteins, and hierarchical clustering was applied to samples. Annotation bars on the top of the heatmap indicate ISG cluster, age, sex, and BMI. The adjacent right panel heatmap summarizes log_2_ fold changes (LFC) for pairwise cluster comparisons (HIS vs LIS, MIS vs LIS, and HIS vs MIS), indicating the direction and magnitude of differential protein abundance between ISG-defined groups. **C)** PCA plot of metabolomic data from HC, Conv, and COVID-19 patients categorized into LIS, MIS, and HIS clusters. The first two principal components are shown, color-coded by cluster categories, highlighting metabolic differences across groups. **D)** UpSet plot depicting the overlap of significantly altered metabolites across ISG-defined clusters (LIS, MIS, HIS). The number of differentially abundant metabolites (LIMMA, FDR < 0.05) in each comparison is indicated, highlighting shared and unique metabolic disruptions across clusters. **E)** Heatmap of significantly altered plasma metabolites across ISG-defined clusters. The main heatmap displays z-score normalized metabolite abundance across individual samples, within LIS, MIS, and HIS clusters. Metabolite abundance is log2 transformed and z-score normalized, with hierarchical clustering applied. Row annotation represents indicated metabolites and their super-pathways, while column annotations indicate individual samples within specific ISG clusters. Annotation bars on the top of the heatmap indicate ISG cluster, age, sex, and BMI. The adjacent right panel heatmap summarizes log_2_ fold changes (LFC) for pairwise cluster comparisons (HIS vs LIS, MIS vs LIS, and HIS vs MIS), indicating the direction and magnitude of metabolic differences between ISG-defined clusters.

To identify specific inflammatory mediators associated with ISG expression, we performed hierarchical clustering using the Olink plasma proteomic data (Figure 3B). Hierarchical clustering revealed distinct inflammatory signatures across ISG clusters, characterized by a progressive increase in inflammatory mediators from LIS to HIS (Figure 3B). Several key inflammatory mediators, including CXCL10, CXCL11, MCP-2, MCP-3, IFN-γ, IL-10, and TNF, showed a graded increase and were significantly elevated in HIS compared to LIS (LIMMA, FDR < 0.05, Table S4), consistent with chemokine-driven inflammatory program associated with interferon signaling. IL-6 and TNF, both implicated in severe inflammatory responses, were particularly upregulated in HIS, suggesting a potential contribution to disease severity within this ISG-defined group.

### Enhanced type I and type II ISG expression in COVID-19 patients reshapes lipid and amino acid metabolism

Increasing evidence supports a close crosstalk between cellular metabolism and IFN signaling during viral infection. Type I IFN has been shown to modulate glycolysis, lipid metabolism, and oxidative phosphorylation in multiple immune cell types [13]. ISGs induced by IFNs also regulate central carbon metabolism [21]. To examine how IFN-driven transcriptional programs relate to systemic metabolism during COVID-19, we analyzed the ISGs cluster-specific changes in global plasma metabolomics data [20]. Principal Component Analysis (PCA) of metabolomics revealed clustering patterns that closely resembled ISG-defined groups (Figure 3C). LIS samples clustered near HC and convalescent samples, whereas MIS and HIS samples showed a progressive divergence, with HIS exhibiting the greatest metabolic shift, indicating increasing metabolic dysregulation with higher ISG expression. UpSet analysis identified the highest number of differentially altered metabolites in the HIS vs. LIS comparison (LIMMA, FDR < 0.05; Table S5), followed by HIS vs. MIS and MIS vs. LIS (Figure 3D), consistent with a graded metabolic shift across ISG clusters.

Lipid metabolism was the most profoundly affected pathway, followed by amino acid metabolism. Multiple lipid species, including several phospholipids, sphingolipids, and carnitines, were significantly reduced in HIS compared to LIS and MIS (Figure 3E), suggesting impaired lipid homeostasis. Amino acids such as arginine, gamma-carboxyglutamate, and 6-bromotryptophan were also reduced in HIS, indicating broader metabolic disruption. Both type I and type II ISG scores showed predominantly a negative association with lipid and amino acid metabolites (Spearman correlation, *p*-value<0.05, Figure S4, Table S6), indicating that heightened interferon-associated inflammation is linked to systemic metabolic depletion. Such metabolic alterations may impair immune cell function, as arginine depletion is associated with T cell dysfunction [22] and tryptophan metabolism plays a key role in immunoregulation [23].

### ISG Expression is Associated with Myeloid Cell Expansion and Enhanced Inflammatory Activation

To relate systemic ISG expression to immune compartment shifts, we inferred immune cell proportions using digital cell quantification (DCQ) based on reference signature of 18 blood cell types from the Human Protein Atlas [24] and the EPIC deconvolution algorithm [25]. Here cell-type estimates were interpreted qualitatively, as relative indicators of immune compartment shifts rather than absolute immune cell frequencies, given the known limitations of whole-blood deconvolution in inflammatory states. DCQ revealed a progressive shift toward an innate immune-dominant state with increasing ISG expression. LIS patients exhibited immune profiles similar to healthy controls, whereas MIS and HIS displayed marked enrichment of innate immune cells, particularly neutrophils and classical monocytes (Figure 4A, Figure S3, Table S7). Targeted phenotyping in a subset of samples supported these findings, showing increased immature neutrophils in HIS and higher classical monocytes in both HIS and MIS (not statistically significant), reinforcing the association between ISG expression and inflammatory myeloid cell expansion (Figure S4).

**Figure 4:**
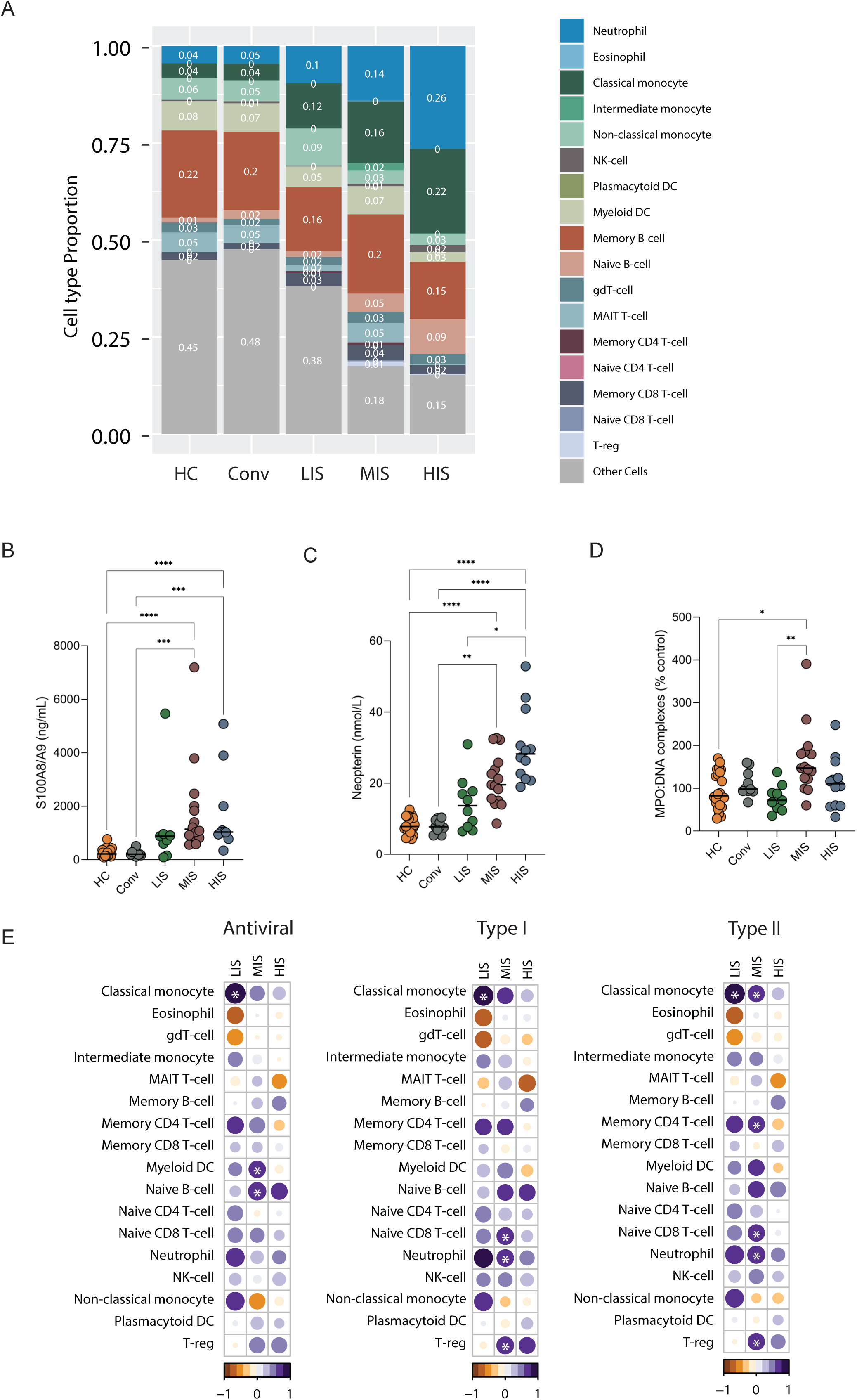
Innate immune activation and association with ISG expression in COVID-19 clusters. **A)** Stacked bar plot showing the proportion of immune cell types inferred from whole-blood transcriptomics data using reference-based Digital Cytometry Quantification (DCQ) across healthy controls (HC), convalescent individuals (Conv), and ISG-defined clusters (LIS, MIS, HIS) reflecting relative differences between groups rather than absolute immune cell frequencies. Cell types with a frequency < 0.05 were grouped under “other cells.” **B-D)** Dot plots depicting plasma levels of key immune activation markers: **(B)** S100A8/A9 (ng/mL), **(C)** neopterin (nMol/mL), and **(D)** MPO:DNA complexes (% of total), indicative of NETosis, across ISG clusters. *p*-values were calculated using the Mann-Whitney U test, with significance denoted as \**p*<0.05, \*\**p*<0.01, \*\*\**p*<0.001, and \*\*\*\**p*<0.0001. **E)** Bubble plot displaying the correlation coefficients between GSVA enrichment scores of inferred immune cell proportions and ISG expression scores (Left-panel: antiviral; Middle-panel: type I IFN; and Right-panel: type II IFN) within each ISG-defined cluster. Circle color represents the direction of correlation (positive or negative Spearman’s rho), while circle size is proportional to the absolute correlation coefficient. Bubbles showing significant correlations (Spearman’s correlation, *p*-value < 0.05) are marked with white asterisks highlighting the association between specific immune cell populations and the interferon response.

To determine whether these expanded innate immune populations were also functionally activated, we measured plasma levels of S100A8/A9 and neopterin, markers of neutrophil and monocyte activation. Both markers were significantly elevated in MIS and HIS compared to LIS (Figure 4B-C; Mann-Whitney U test, *p*-value < 0.05, Table S8), indicating enhanced myeloid activation with increasing ISG expression.

Neutrophil activation can also lead to the formation of neutrophil extracellular traps (NETs), which interact with cytosolic DNA sensors such as cGAS and activate IFN pathways via cGAS-STING signaling [26]. To assess NET formation, we measured MPO:DNA complexes. MPO:DNA levels were significantly increased in MIS, but not in HIS (Figure 4D), indicating that NETosis is not a defining feature of the ISG-high phenotype. Stratification by disease severity showed that S100A8/A9 and neopterin were elevated in both mild and severe cases, whereas MPO:DNA levels did not differ by severity (Figure S5). These findings indicate that NET-associated inflammation represents a distinct immune activation program that is separable from the ISG-high endotype and does not drive severity within the HIS group.

Correlation analysis revealed that LIS showed positive associations with classical monocytes across all ISG pathways, while MIS correlated with additional immune subsets (Figure 4E; Spearman correlation, FDR < 0.05, Table S9). Notably, no immune cell type showed a significant positive correlation with ISG scores within HIS, suggesting a more complex immune configuration rather than expansion of discrete cell populations.

Together, these findings indicate that high ISG expression is associated with a systemic innate immune-activated state characterized by relative myeloid enrichment and elevated inflammatory mediators. However, the absence of strong correlations between ISG scores and specific immune subsets within HIS suggests immune dysregulation rather than simple cell expansion. Because HIS includes both mild and severe cases, ISG expression alone is insufficient to explain disease severity, motivating further investigation into additional modulatory factors.

### Metabolic and Immune Dysregulation Define Disease Severity in the High ISG-expressing Group by altering Neutrophil and Monocyte Activation

Because severe disease was observed in both LIS and HIS, we focused on the HIS group to identify determinants of severity within a shared high-ISG context. This approach allowed us to interrogate factors associated with disease severity beyond the presence of a strong systemic interferon-stimulated gene signature. Digital cell quantification (DCQ) analysis showed that HIS patients had increased neutrophil and monocytes proportions and elevated S100A8/A9 and neopterin levels, indicating that innate immune activation is a defining feature of HIS. However, disease severity was not associated with further increases in myeloid expansion. Moreover, no immune cell type correlated positively with ISG scores within HIS, and NETosis was not elevated, suggesting that functional immune alterations and systemic inflammatory or metabolic factors may underlie severity within this group.

To assess immune activation within the HIS endotype, we incubated healthy isolated neutrophils and PBMCs (for monocytes) with plasma from mild and severe cases and measured activation markers by flow cytometry. Functional assays were restricted to plasma from HIS patients to isolate severity-associated effects within a shared high-ISG background and avoid confounding by differences in baseline interferon responsiveness across ISG clusters. Neutrophils activation was assessed using CD66b, CD11b, and CD62L, while monocyte activation was evaluated using CD169 (Siglec-1). Interestingly, plasma from severe cases showed a tendency towards reduced neutrophil activation, characterized by lower CD66b and CD11b expression, and preserved CD62L expression, together with significantly reduced monocyte activation (low CD169; *p*<0.05) compared to plasma from mild cases (Figure 5A). Though we observed a clear trend, substantial inter-individual variability was observed, and not all comparisons reached statistical significance. These findings indicate a trend toward functional suppression of innate immune activation by plasma from severe HIS cases, while highlighting heterogeneity within this group.

**Figure 5:**
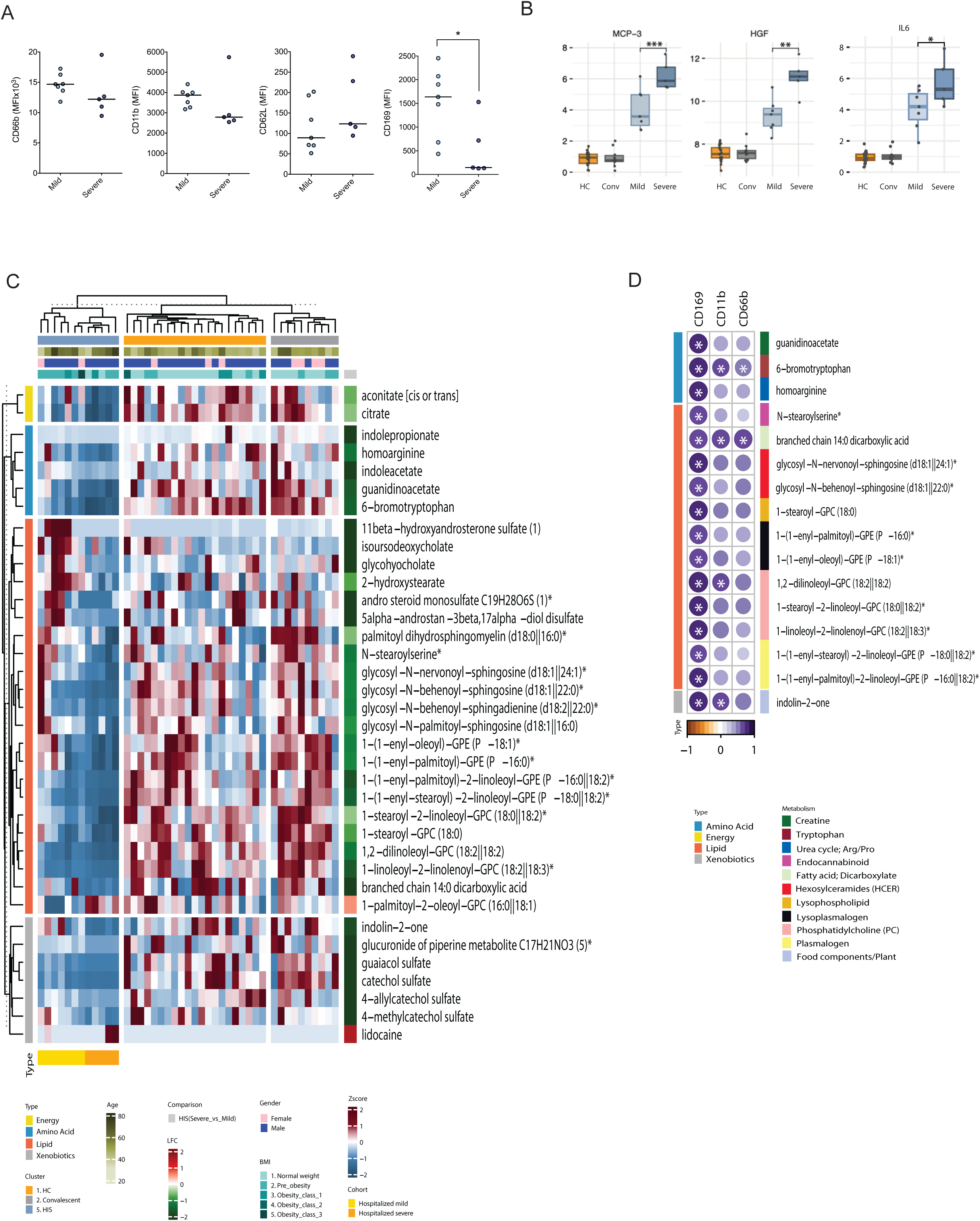
Immune and metabolic determinants of severity in the HIS cluster. **A)** Dot plots showing the differential ability of plasma from mild and severe HIS cases to activate healthy neutrophils and monocytes in an ex vivo assay. Activation was assessed using CD66b/CD11b for neutrophils and CD169 for monocytes. Lower activation in severe cases suggests the presence of suppressive soluble factors in circulation. Plasma used in the functional assays was derived exclusively from HIS patients and stratified into mild HIS and severe HIS. Healthy donor neutrophils and PBMCs were used as responder cells. **B)** Boxplots showing the plasma levels of key inflammatory mediators HGF, MCP-3 (CCL7), and IL-6, in mild (light blue) and severe (dark blue) HIS patients. Severe HIS cases exhibited significantly higher levels of MCP-3, HGF, and IL-6 compared to mild HIS cases. Adjusted *p*-values were calculated using LIMMA, with significance thresholds: * *p*<0.05, ** *p*<0.01 and *** *p*<0.001. **C)** Heatmap of log₂-scaled and Z-score-transformed metabolite levels significantly altered between mild and severe HIS cases. Xenobiotics were excluded. Rows represent super-pathways of metabolites, and columns represent individual HIS patient samples. **D)** Bubble plot showing Spearman correlation coefficients between significantly altered metabolites and neutrophil (CD66b, CD11b) and monocyte (CD169) activation markers in HIS patients. *p*-values < 0.05 were considered statistically significant. Bubble color represents the direction and magnitude of the correlation coefficient, as indicated by the color scale. Bubble size reflects the absolute strength of the correlation (Spearman’s rho) with larger bubbles indicating stronger associations. Asterisks denote statistically significant correlations (*p-*value < 0.05, Spearman’s correlation). Only metabolites significantly altered in HIS compared to LIS or MIS (LIMMA, FDR < 0.05) were included.

To identify soluble mediators associated with disease severity in HIS, we compared inflammatory factors (Olink onco-Immunology panel), between severe and mild cases using LIMMA (Table S10). MCP-3, HGF, and IL-6 were significantly elevated in severe HIS cases (Figure 5B, FDR < 0.05). MCP-3 (CCL7) is a key monocyte-recruiting chemokine [27], consistent with monocyte involvement in severe disease, while HGF, a regulator of tissue repair and immune modulation [28], and IL-6, a major pro-inflammatory cytokine [29], have been linked to immune dysregulation and adverse COVID-19 outcomes. These results indicate that severity within HIS is associated with dysregulated inflammatory signaling rather than ISG magnitude alone.

Metabolomic profiling identified 36 significantly altered metabolites between mild and severe cases (Figure 5C, LIMMA FDR < 0.05, Table S11), with energy and lipid metabolism being the most dysregulated pathways. TCA cycle intermediates, particularly citrate and aconitate, were significantly reduced in severe cases, suggesting impaired mitochondrial metabolism which is essential for proper immune activation [30]. Lipid classes such as plasmalogens, phosphatidylcholines, and sphingolipids were also significantly disrupted, potentially affecting membrane integrity, immune signaling, and cell survival [31].

Correlation analysis between significantly altered metabolites and neutrophil/monocyte activation markers (FACS MFI) revealed significant associations for 16 out of 36 significantly altered metabolites with at least one immune activation marker (Figure 5D; Table S12, Spearman correlation, *p*-value<0.05). Several metabolites, including 6-bromotryptophan and branched-chain 14:0 dicarboxylic acid, correlated with both neutrophil (CD11b, CD66b) and monocyte (CD169) activation markers, while others showed stronger associations with monocyte activation. 1,2-dilinoleoyl-GPC (18:2II18:2) correlated with both CD169 and CD11b. Consistent with these findings, key severity-associated inflammatory mediators (MCP-3, HGF, and IL-6) showed predominantly negative correlation trends with multiple metabolites (Figure S6), suggesting that heightened inflammatory signaling coincides with systemic metabolic depletion. These findings suggest that metabolic alterations, particularly in tryptophan metabolism, lipid oxidation, and phospholipid remodeling, may influence immune activation and contribute to disease severity within HIS.

In an exploratory analysis of a small subset of patients with available PBMCs, we assessed ISG protein expression by western blot and compared it to transcript levels. In mild cases, ISG15, RIG-I, IRF3, and STING protein levels correlated positively (Spearman’s Rho > 0.6) with their corresponding transcripts, whereas this relationship was lost in severe cases (Figure S7). Although limited by sample size, this observation raises the possibility of post-transcriptional or translational dysregulation contributing to impaired antiviral responses in severe disease.

## Discussion

The interferon (IFN)-signaling is activated immediately upon viral infection and plays a pivotal role in antiviral immunity. A balanced IFN response is crucial to eliminating viral infections while preventing excessive inflammation and immune-mediated damage. While timely IFN responses promote viral clearance and are associated with milder COVID-19 outcomes [4, 32], dysregulated or delayed IFN signaling has been implicated in severe disease [8, 9] [33–35]. In particular, excessive or prolonged IFN-associated inflammation, often described as interferonopathy has been linked to immune-mediated tissue damage in critical COVID-19 [36–38]. Together, these observations highlight the dual nature of IFN responses [39, 40] and highlight the need to define when IFN-driven programs are protective versus pathogenic in human disease.

In this study, we identify distinct systemic interferon-associated immune-metabolic endotypes in COVID-19 based on whole-blood interferon-stimulated gene (ISG) expression. While high ISG expression reflects a state of heightened interferon-associated immune activation, our data demonstrate that disease severity emerges only when this transcriptional state coincides with specific inflammatory mediators and metabolic constraints that impair effective innate immune function. This distinction between ISG magnitude and immune functionality provides a framework to reconcile previously conflicting reports on IFN responses in COVID-19 and emphasizes that ISG expression alone is insufficient to predict clinical outcome.

We observed pronounced heterogeneity in systemic ISG expression among COVID-19 patients that was largely independent of disease severity. Severe COVID-19 patients exhibited higher antiviral ISG scores than healthy controls (Figure 1C), confirming intact induction of interferon-driven antiviral programs at the systemic level. However, the occurrence of severe disease despite robust antiviral ISG expression indicates that IFN-driven transcriptional activation alone does not guarantee protective immunity. Stratification based on whole-blood ISG expression revealed three distinct clusters: low ISG score (LIS), moderate ISG score (MIS), and high ISG score (HIS), each characterized by distinct immune activation state, inflammatory profiles, and metabolic reprogramming. This classification highlights the complex interplay between IFN signaling, innate immune activation, and systemic metabolism during acute SARS-CoV-2 infection.

The distribution of clinical severity across these ISG-defined endotypes further supports a non-linear relationship between interferon responses and disease outcome. Mild cases were enriched within the MIS and HIS groups, whereas severe disease occurred predominantly at the extremes, within both LIS and HIS, with relatively few severe cases in MIS (Figure 2B). This pattern suggests that intermediate systemic ISG activation may reflect a balanced immune state compatible with antiviral control, while severe disease may arise either in the context of insufficient interferon activity (LIS) or sustained interferon-associated inflammation (HIS). Supporting this interpretation, principal component analyses of plasma metabolomics and proteomics restricted to severe cases revealed that severe HIS patients segregated from severe LIS and MIS patients despite comparable clinical severity (Figure S8), indicating that severe COVID-19 can arise through biologically distinct immune-metabolic trajectories rather than a single severity continuum.

Analysis of circulating interferon transcripts and proteins further clarifies the relationship between interferon signaling and systemic ISG expression. Although low-level IFN mRNA transcripts were detectable in whole-blood RNA sequencing, plasma concentrations of type I and type III interferons did not differ consistently across disease severity groups and showed no correlation with ISG scores (Figure 1D). This uncoupling contrasts with influenza infection, where interferon protein levels closely track antiviral gene expression [41], and is consistent with reports describing temporally dysregulated interferon responses in COVID-19 [10]. Together, these findings indicate that IFN mRNA and protein measurements provide limited explanatory power for systemic ISG heterogeneity and support the interpretation that ISG signatures reflect downstream, context-dependent interferon-associated immune states rather than contemporaneous interferon abundance.

Neutralizing autoantibodies against type I interferons have been implicated in severe COVID-19 by impairing IFNAR signaling and downstream ISG activation [42, 43]. In our cohort, IFN antigen reactivity was detected in a small subset of patients across ISG-defined clusters using a stringent cutoff compared with other studies [44, 45], with functional neutralization observed in several cases. However, IFN antigen reactivity did not correlate with disease severity, ISG expression levels, or circulating interferon protein abundance. These findings indicate that IFN antigen reactivity does not account for the systemic interferon-associated immune-metabolic endotypes identified in this study and is unlikely to be a primary driver of ISG heterogeneity in this cohort. The detection of IFN antigen reactivity in convalescent plasma further raises questions about the persistence of immune dysregulation beyond acute infection, but larger longitudinal studies will be required to define its clinical relevance.

A defining feature of the HIS endotype was enrichment of innate immune activation, reflected by increased neutrophil and monocyte proportions inferred by DCQ and elevated plasma levels of S100A8/A9 and neopterin, markers of neutrophil and macrophage activation, respectively. Neutrophils are key effectors of innate immunity and express cytosolic pattern-recognition receptors such as RIG-I and MDA5, enabling interferon-associated activation upon viral sensing [46]. Upon activation, neutrophils can release S100A8/A9, an alarmin that functions as a damage-associated molecular pattern to amplify inflammatory signaling and recruit additional immune cells [47]. S100A8/A9 has been implicated in SARS-CoV-2–associated immunopathology [48], and its elevation in MIS and HIS in our cohort supports its role as a marker of interferon-associated myeloid activation rather than a direct determinant of disease severity. Consistent with this, SARS-CoV-2 infection has been shown to induce alarmin release, promoting expansion and activation of immature neutrophils and contributing to dysregulated innate immune responses [49]. Neopterin, released by activated macrophages upon IFN-γ induction [50], followed a similar pattern, further supporting the role of IFN-driven myeloid cell activation within the HIS cluster. ISGs can be induced through both IFN-dependent JAK-STAT signaling and IFN-independent IRF3 activation [51, 52]. In COVID-19, neutrophil activation has been linked to NET formation, [53] which can further amplify interferon signaling via cGAS-STING activation [26]. However, despite previous reports linking NETs to severe COVID-19 [54], our findings did not indicate any association between NET formation and either disease severity or HIS endotype. Instead, NETosis was most prominent in the MIS group, indicating that NET-associated inflammation represents a distinct immune activation program that is separable from the ISG-high systemic state.

Within the HIS endotype, disease severity was associated with a coordinated inflammatory and metabolic state rather than with the magnitude of interferon-stimulated gene expression alone. Severe HIS cases exhibited elevated circulating levels of MCP-3 (CCL7), hepatocyte growth factor (HGF), and IL-6, mediators implicated in monocyte recruitment [27], tissue remodeling, and systemic immune dysregulation in COVID-19 [28, 29]. These inflammatory signals coincided with depletion of tricarboxylic acid (TCA) cycle intermediates, including citrate and cis-aconitate, and reduced abundance of multiple lipid classes involved in membrane integrity and immunometabolic signaling. Such metabolic perturbations are consistent with impaired mitochondrial function and reduced bioenergetic capacity, conditions known to limit effective innate immune cell activation.

During viral infection, immune cells undergo extensive metabolic reprogramming to meet the energetic and biosynthetic demands of innate immune activation, while viruses can exploit host metabolic pathways to support its own replication [55] [56]. In line with this, our metabolomic profiling revealed significant alterations in amino acid and lipid metabolism, with suppression of TCA cycle emerging as a defining feature of severe disease in HIS endotype. Reduced circulating levels of citrate and aconitate suggest mitochondrial dysfunction, which may impair energy metabolism and immune cell function [30]. In parallel, depletion of lipids such as plasmalogens, phosphatidylcholines, and sphingolipids which are essential for membrane stability, receptor signaling and immune cell communication [31, 57] is consistent with impaired immune signaling and defective inflammatory resolution. These findings are concordant with reports from independent COVID-19 cohorts across different geographical regions, which have described broad perturbations in amino acid and lipid metabolisms in severe or fatal COVID-19 [58]. However, unlike prior studies, our analysis directly links these metabolic alterations to interferon-associated immune states, highlighting a previously underexplored connection between IFN-driven inflammation and systemic metabolic dysfunction.

Experimental models have shown that reduced lipid biosynthesis can itself induce type I IFN responses in a STING dependent manner [59], providing a mechanistic framework that falls in line with our observation of lowered lipid metabolite levels in patients with high ISG expression. Importantly lipid depletion was further associated with the disease severity in both HIS and MIS clusters, with distinct lipid signatures observed in each group, suggesting that interferon-associated metabolic stress represents a broader feature of COVID-19 immunopathology rather than a uniform consequence of disease severity. Notably, reduced levels of citrate and cis-aconitase, central intermediates linking the TCA cycle to fatty acid metabolism were unique to severe HIS cases. Both metabolites support the production of itaconate, a metabolite with well established anti-inflammatory and IFN-regulatory function and its derivatives have inhibitory effects on virus replication, including SARS-CoV-2 [60–62]. Their depletion suggests loss of a key immunoregulatory metabolic axis in severe disease. Correlation analyses linking these metabolic alterations to reduced neutrophil (CD66b, CD11b) and monocyte (CD169) activation further support a model in which metabolic constraints actively limit immune cell responsiveness rather than simply reflecting downstream consequences of inflammation.

Sex is a recognized risk factor for severe COVID-19, with male patients exhibiting higher rates of hospitalization and adverse outcomes [63, 64]. In our cohort, all severe cases were male, limiting our ability to disentangle sex-specific effects from disease severity. However, comparison of ISG scores between male and female patients revealed no significant differences across antiviral, type I IFN-associated, or type II IFN-associated ISG signatures (Figure S9). These findings suggest that sex-associated risk is unlikely to be mediated by differences in systemic ISG induction and may instead reflect downstream differences in immune regulation, metabolic resilience, or tissue-specific immune responses not captured here. Larger, sex-balanced cohorts will be required to resolve these mechanisms.

Our study has several limitations. First, metabolic alterations were assessed in plasma, precluding direct conclusions about intracellular metabolic changes in immune cells. Second, while significant associations between metabolic changes and immune activation were observed, causal relationships remain to be established. Third, reliance on whole-blood bulk transcriptomics limits assignment of ISG expression to specific cell types, and digital cell quantification was therefore interpreted qualitatively. Fourth, COVID-19 immunopathology is highly compartmentalized, and peripheral blood analyses may not fully capture tissue-resident immune responses in organs such as the lung or kidney. Our analyses are limited to peripheral blood defining systemic immune-metabolic states reflected in circulation and therefore do not directly capture tissue-resident immune cell programs or local antiviral responses. Finally, the absence of disease or vaccination controls and the modest cohort size limit generalizability. Although this cohort derives from the first pandemic wave, the host response pathways interrogated here including interferon signaling, innate immune activation, and immunometabolic regulation represent fundamental biological processes that are unlikely to be variant-specific. External validation of these immune-metabolic endotypes is currently constrained by the lack of harmonized patient-level datasets integrating bulk transcriptomics, broad plasma metabolomics, and immune activation phenotyping with comparable severity metadata.

In summary, our findings demonstrate that systemic ISG expression delineates distinct immune-metabolic endotypes in COVID-19, but ISG magnitude alone does not determine disease severity. Instead, severity emerges from the convergence of interferon-associated inflammation with metabolic constraints that impair effective innate immune function. These results position immunometabolism as a central determinant of disease outcome and suggest that therapeutic strategies aimed at restoring metabolic homeostasis such as modulation of lipid metabolism or replenishment of TCA cycle intermediates may enhance immune competence in selected patient subsets. More broadly, understanding how IFN-driven immune programs intersect with metabolism may inform precision approaches to viral infections and inflammatory diseases beyond COVID-19.

## Methods

### Study design and patients

The study included both PBMCs and plasma from SARS-CoV-2-infected individuals with available whole-blood transcriptomics data (*n=37*) hospitalized in Stockholm, Sweden during May 2020. These individuals were separated into the category of mild (*n=26*) or severe (*n=11*) disease based on O_2_ requirement (O_2_ consumption <4 L/min and >4 L/min for mild and severe respectively). The cohort was further complemented with SARS-CoV-2 negative individuals (*n=31*), comprising convalescent individuals after SARS-CoV-2 infection (*n=10*) and non-infected healthy controls (*n=21*). The clinical and demographic parameters of the cohort are as earlier described [18, 20]. The study was approved by the Swedish Ethical Review Authority (dnr 2020-01865). All individuals gave informed consent, and the study was conducted according to the Declaration of Helsinki.

### Measurement of plasma IFN levels

Plasma samples were used to measure the levels of IFNs using LEGENDplex Human type 1/2/3 interferon panel (BioLegend #740396), measuring concentrations of IFN-ɑ2, β, 𝛾, 𝜆1, and 𝜆2/3. The assay was performed according to the manufacturers protocol. Samples were acquired on Fortessa (BD Biosciences), and analysis performed in FlowJo 10.8.1 (TreeStar Inc). For analysis samples below the background detection were set to 0 and results reported as the median MFI.

### Measurement of plasma cellular activation markers

Plasma levels of neopterin (Tecan, Cat# RE59321) and S100A8/A9 (Alarmin/Calprotectin) in plasma were measured using enzyme-linked immunosorbent assay (DuoSet ELISA, R&D Systems, USA; cat number DY8226-05), according to the manufacturer’s instructions. Circulating NETs were measured using an MPO-DNA ELISA, with slight modifications from previous descriptions [54, 65]. An anti-MPO antibody (4 µg/mL; clone 4A4, Bio-Rad Laboratories, USA) was used as the capture antibody and an anti-dsDNA antibody from the Cell Death Detection ELISA kit (Roche Diagnostic, Switzerland) was used for detection. Plasma was used at 1:10 dilution in 1% BSA/PBS and incubated for 2 hours at room temperature. The development was done with TMB (BD Biosciences, USA), and the reaction stopped with a 2N sulfuric acid solution. Absorbance was measured at 450 nm and wavelength correction at 540 nm to subtract background using a microplate reader (SpectraMAX 340, USA).

### SARS-CoV-2 antibody and IFN antigen-reactivity screening

Plasma samples were analyzed for antibody reactivity against a panel of recombinant protein antigens using a broad multiplex bead-based array, as described previously [66]. The antigen panel included type I, type II, and type III interferons/cytokines: IFNA1, IFNA2, IFNA4, IFNA5, IFNA6, IFNA7, IFNA8, IFNA10, IFNA14, IFNA16, IFNA17, IFNA21, IFNB1, IFNE, IFNG, IFNK, IFNL4, IFNW1, IL28A, IL28B, IL29, GM-CSF, IL1F6, IL1RN, IL12, IL17A, IL17F, IL22, IL23, and ISG15. Antibody reactivity against SARS-CoV-2 spike, receptor-binding domain (RBD), nucleocapsid, and Epstein-Barr virus EBNA1 was also assessed. Briefly, recombinant proteins were coupled to magnetic beads (MagPlex®, Luminex Corp., Austin, Texas) using AnteoTech Activation Kit for Multiplex Microspheres (Cat# A-LMPAKMM-10) with commercial versions of the proteins (1.5 x 10^6^ beads were coupled using 3 µg of proteins). One microliter of sample was diluted 1:25 in PBS and later diluted 1:10 in 0.05% PBS with 3% BSA and 5% Milk. The sample dilution was subsequently incubated for two hours with 5 µL of the bead solution on a 650 rpm shaker. After three washings in 0.05% PBS, beads were resuspended in 50 µL of 0.2% PFA during 10 minutes. The secondary antibody (Invitrogen, H10104 lot#2384336) incubation lasted for 30 minutes prior to the final analysis in FlexMap 3D instrument (Luminex Corp). For each sample-antigen pair, antibody binding was quantified as mean fluorescence intensity (MFI), calculated from multiple beads coupled to the same antigen. A cutoff for positivity was defined for each antigen as the mean MFI of the healthy control group + 7 SD.

### Neutralization Analysis for Type I Interferon Autoantibodies

Neutralization assays were performed on samples identified for further analysis based on high ISG scores or elevated responses to type I interferons in a multiplex screening assay (threshold: mean + 7 SDs). The assay followed previously reported cell culture-based neutralization protocols [67], utilizing a dual-luciferase reporter system in HEK293T cells to assess the functional inhibition of interferon signaling. On day 1, HEK293T cells in logarithmic growth were detached using 0.01% trypsin, washed, and resuspended at 1×10⁶ cells/mL in DMEM (Gibco) supplemented with 10% FBS and 100 U/mL penicillin-streptomycin. A total of 35,000 cells per well were plated in a 96-well flat-bottom plate. After 3-4 hours of incubation at 37°C to allow for attachment, cells were transfected using Firefly pGL4.45[luc2P/ISRE/Hygro] (IFN-stimulable) and Renilla pRL-SV40 (constitutive control) vectors (Promega) with X-tremeGENE9 transfection reagent (Sigma-Aldrich) at a 3:1 ratio (µL:µg) in OptiMEM medium. The final transfection mix contained 0.8 µg Firefly and 0.4 µg Renilla plasmid per well. Transfection controls were included, and plates were incubated at 37°C for 24 hours. On day 2, stimulation was initiated by adding 5 µL of IFN-α2 (100 ng/mL, MedChemExpress) to achieve a final concentration of 10 ng/mL per well, followed immediately by 5 µL of plasma samples (1:10 final dilution in a total 50 µL volume). Non-stimulation controls received media instead of IFN-α2, and non-plasma technical controls received media instead of plasma. Positive control samples from APS1 patients (known neutralizing autoantibodies) and negative control blood donor plasma were included. On day 3, following 24 hours of incubation, neutralization was assessed using the Dual-Luciferase Reporter Assay System (Promega). Cells were lysed, and luminescence was measured using a plate reader (Tecan, Magellan) with Firefly (magenta filter) and Renilla (green filter) detection. The Firefly-to-Renilla ratio was calculated, with Renilla readings serving as internal controls to correct for variations in transfection efficiency and cell viability. APS1 patient samples exhibited complete neutralization (>99%, Firefly/Renilla ratio <0.050), while healthy donor samples showed no neutralization (>2.000 ratio). A standard response curve was generated using APS1 samples (0% response) and blood donors (100% response). Definite neutralization was defined as >90% response reduction, while partial neutralization was categorized as 80–90% response reduction. The assay controls performed as expected, confirming the robustness of the neutralization assessment.

### Data and preprocessing

Count tables for transcriptomics, olink proteomics data, and metabolomics data were retrieved and reanalyzed from Ambikan et al. 2022 [18]. Before analysis, transcriptomics data were filtered for low variance (variance < 0.2) and transformed using variance-stabilizing transformation (VST) from R package DESeq2 [68]. Genes expression profile associated with the Reactome (https://reactome.org) pathways antiviral mechanisms, interferon alpha/beta and interferon gamma were extracted from the data. ISG scores were calculated for each patient by making the sum of Z-scaled expression values of genes linked to each pathway [19]. Plasma metabolomics data were derived from a previously published dataset generated from the same cohort and reanalyzed here in the context of ISG-defined stratification [20]. Metabolomics data were log_2_-transformed before analysis. The data were obtained from standardized metabolomics platform which enables consistent metabolite annotation and cross-sample comparison but limits direct feature-level matching with discovery-based mass spectrometry datasets commonly used in public COVID-19 cohorts.

## Statistics

Spearman pairwise correlations between features were done using the R package Psych. R package limma for differential abundance analysis was used in metabolomics and proteomics data [69] with adjustment for age, sex and BMI category. R package DESeq2 was used to perform differential gene expression analysis in transcriptomics data [68]. Factors of possible unwanted variation in the data estimated using R package RUVSeq [70] were adjusted along with age, gender and BMI category while executing DESeq2. Means of continuous variables were compared using Mann-Whitney U test. All *p-values* were adjusted using false discovery rate (FDR) and by default FDR < 0.05 was considered significant. Different cutoffs are indicated in the analysis. Absolute fold change higher than 1 was used as a supplementary cutoff in transcriptomics data.

## Cell profiling using digital cell quantification (DCQ)

Digital cell quantification (DCQ) was performed to estimate immune cell proportions from bulk RNA sequencing data using the EPIC deconvolution algorithm [25] with a reference signature of 18 blood cell types from the Human Protein Atlas [24]. Transcripts per million (TPM) normalized gene expression data was used for the analysis. The deconvolution approach utilized a constrained least-squares regression to infer relative immune cell proportions based on predefined reference profiles. To assess the relationship between interferon-stimulated gene (ISG) expression and immune cell distributions, DCQ-derived cell fractions were correlated with ISG scores using Spearman’s rank correlation with false discovery rate (FDR) correction for multiple comparisons. All statistical analyses were conducted using R (v4.1.0), with significance thresholds set at FDR < 0.05 unless otherwise specified.

## Clustering

Patient clustering was performed based on interferon-stimulated gene profiles from transcriptomics data using R package ConsensusClusterPlus. From 2 to 10 clusters were tested (maxK) on 1000 subsamples (reps), with 0.8 of items (pItem) and all features (pFeature) to sample. The clustering algorithm used was hierarchical clustering (hclust). The number of clusters was determined based on Consensus Cumulative Distribution Function (CDF) and delta area plots.

## Visualisation

Scatter plots, box plots, bubble plots, dot plots, bar plots, PCA plots and volcano plots were done using R packages ggplot2 and ggpubr. Heatmaps were done using R package ComplexHeatmap.

## Plasma Treatment of Neutrophils and PBMCs for Monocyte Activation and Flow Cytometry Analysis

Neutrophils and peripheral blood mononuclear cells (PBMCs) were isolated from buffy-coated blood of healthy donors using density gradient centrifugation with Ficoll-Paque PLUS (GE Healthcare). Whole blood was diluted 1:1 with phosphate-buffered saline (PBS), layered over Ficoll, and centrifuged at 400 × g for 30 min at room temperature without brake. The PBMC layer was collected, washed twice in PBS, and resuspended in RPMI-1640 (Gibco) supplemented with 2 mM L-glutamine and 1% penicillin-streptomycin. Neutrophils were isolated from the erythrocyte pellet via hypotonic lysis, followed by washing and resuspension in RPMI-1640 with 2 mM L-glutamine and 1% penicillin-streptomycin.

PBMCs were seeded in 96-well U-bottom plates at 2 × 10^5^ cells/well, and neutrophils were plated in 96-well flat-bottom plates at 2 × 10^5^ cells/well. Cells were treated with 10% plasma (1:10 dilution in culture medium) from COVID-19 patients stratified by disease severity (mild vs. severe) within the high ISG expression (HIS) group. Control wells received plasma from healthy blood donors (negative control). PBMCs were incubated for 24 hours and neutrophils for 4 hours at 37°C with 5% CO₂.

Following incubation, cells were washed and stained with fluorophore-conjugated antibodies for activation markers. PBMCs were stained for monocyte activation using CD169 (Siglec-1, clone 7-239, BV421; BD Biosciences), a marker induced by type I IFNs, along with CD14 (clone M5E2, PerCP; BioLegend) and HLA-DR (clone L243, PE/Cy7; BioLegend) for monocyte gating. Neutrophil activation was assessed using CD11b (clone ICRF44, PE; BioLegend), CD66b (clone G10F5, FITC; BioLegend) and CD62L (clone DREG-56, BV421; BioLegend). A viability dye (Live/Dead Fixable, Thermo Fisher) was used to exclude dead cells. Samples were acquired on a BD FACSVerse flow cytometer, and data were analyzed using FlowJo v10.

Mean fluorescence intensity (MFI) values for CD169, CD11b, CD66b, and CD62L were compared between treatment groups using Kruskal-Wallis tests followed by Dunn’s post-hoc analysis. Correlations between metabolic alterations and immune activation markers were assessed using Spearman’s rank correlation with false discovery rate (FDR) correction. Statistical analyses were conducted in GraphPad Prism 9 and R v4.1.0, with significance set at FDR < 0.05 unless otherwise specified.

## Acknowledgements

The authors would like to thank the excellent support received from the study nurses Elisabet Storgard and Ronnie Ask, Södersjukhuset. We thank the National Facility for Autoimmunity and Serology Profiling at SciLifeLab for excellent technical support related to the antibody-reactivity screening experiments. We also acknowledge Magdalini Lourda, Karolinska Institutet for meaningful discussions that contributed to this work.

## Authors Contribution

S.G. Conceptualized the project; S.G., U.N., N.L. and R.L-J., acquired the funding; R.L-J., A.C., S.R., A.Y., S.S.A, X.C., M.S. and M.A.-G., performed research; A.A. and F.M., performed bioinformatic analysis; H.N. and C.J.T performed clinical research. N.L. and U.N. contributed new reagents/analytic tools; A.A., R.L-J., S.R., A.C., A.Y., F.M., S.S.A, N.L., U.N. and S.G. analyzed data; S.R., A.A., A.Y., R.L-J., A.C., F.M., S.S.A and S.G. wrote the original draft of the paper. A.C., R.L-J., S.R., X.C., M.S., H.N., C.J.T, N.L. and U.N. reviewed and edited the manuscript. All authors have reviewed and approved of the final version.

