## Supplemental Figures for "Systemic Multi-Omics Analysis Reveals Interferon Response Heterogeneity and Links Lipid Metabolism to Immune Alterations in Severe COVID-19"

**Figures Supplementary:**

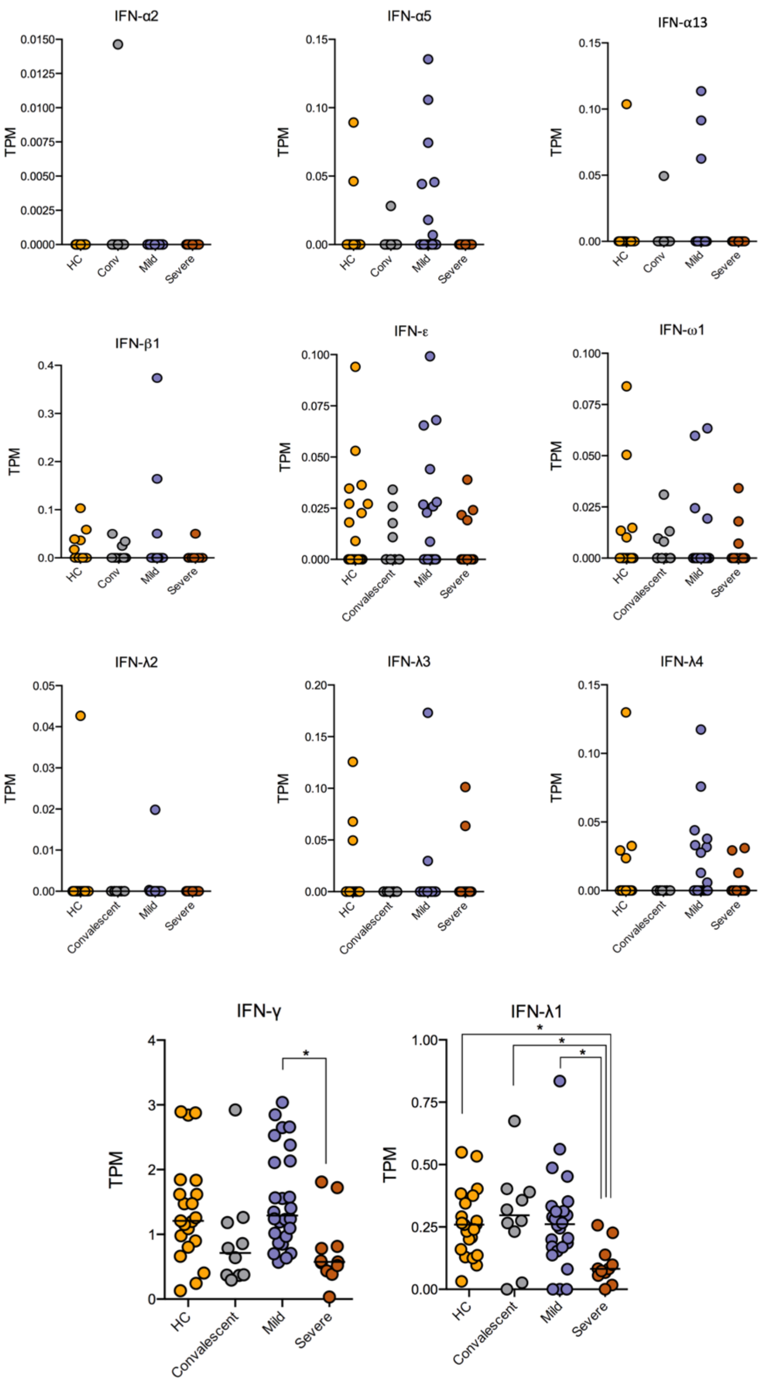


**Supplementary Figure S1:** Transcriptomic levels of detectable type-I, type-II, and type-III interferon (IFN) mRNA expression across different clinical groups. TPM (transcripts per million) values are displayed for individual IFN subtypes, including type-I IFNs (IFN-α2, IFN-α5, IFN-α13, IFN-β1, IFN-ε, and IFN-ω1), type-II IFN (IFN-γ), and type-III IFNs (IFN-λ2, IFN-λ3, IFN-λ4, and IFN-λ1). Data are stratified by healthy controls (HC), convalescent individuals, and COVID-19 patients with mild and severe disease. Statistical significance between groups was assessed using the Kruskal-Wallis test, with post hoc comparisons indicated by * $p<0.05$.

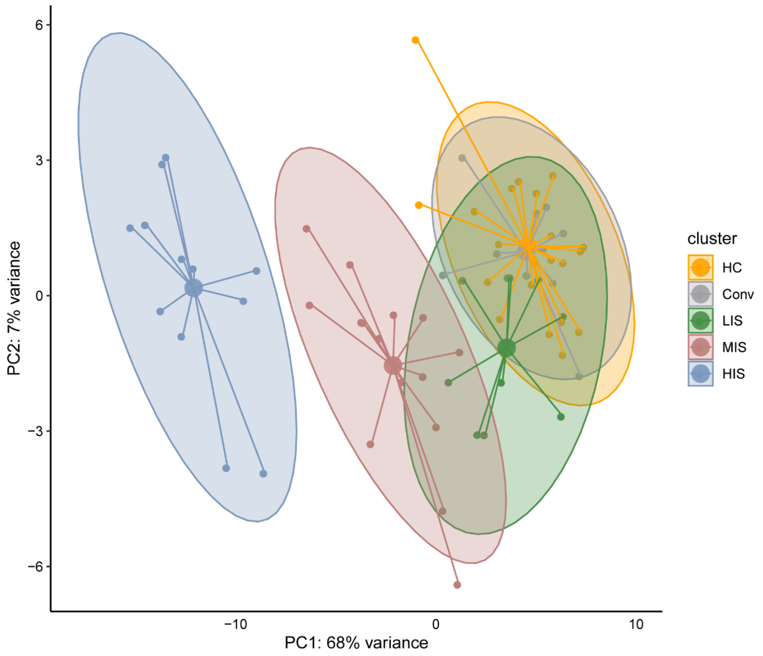


**Supplementary Figure S2:** Principal component analysis (PCA) plot of whole-blood transcriptomic data, illustrating the clustering of study participants based on ISG-defined groups. Clusters include HC (healthy controls), Conv (convalescent), LIS (low ISG score), MIS (moderate ISG score), and HIS (high ISG score). Each dot represents an individual sample, and ellipses indicate the 95% confidence interval for each cluster. Principal component 1 (PC1) and principal component 2 (PC2) explain 68% and 7% of the variance, respectively.

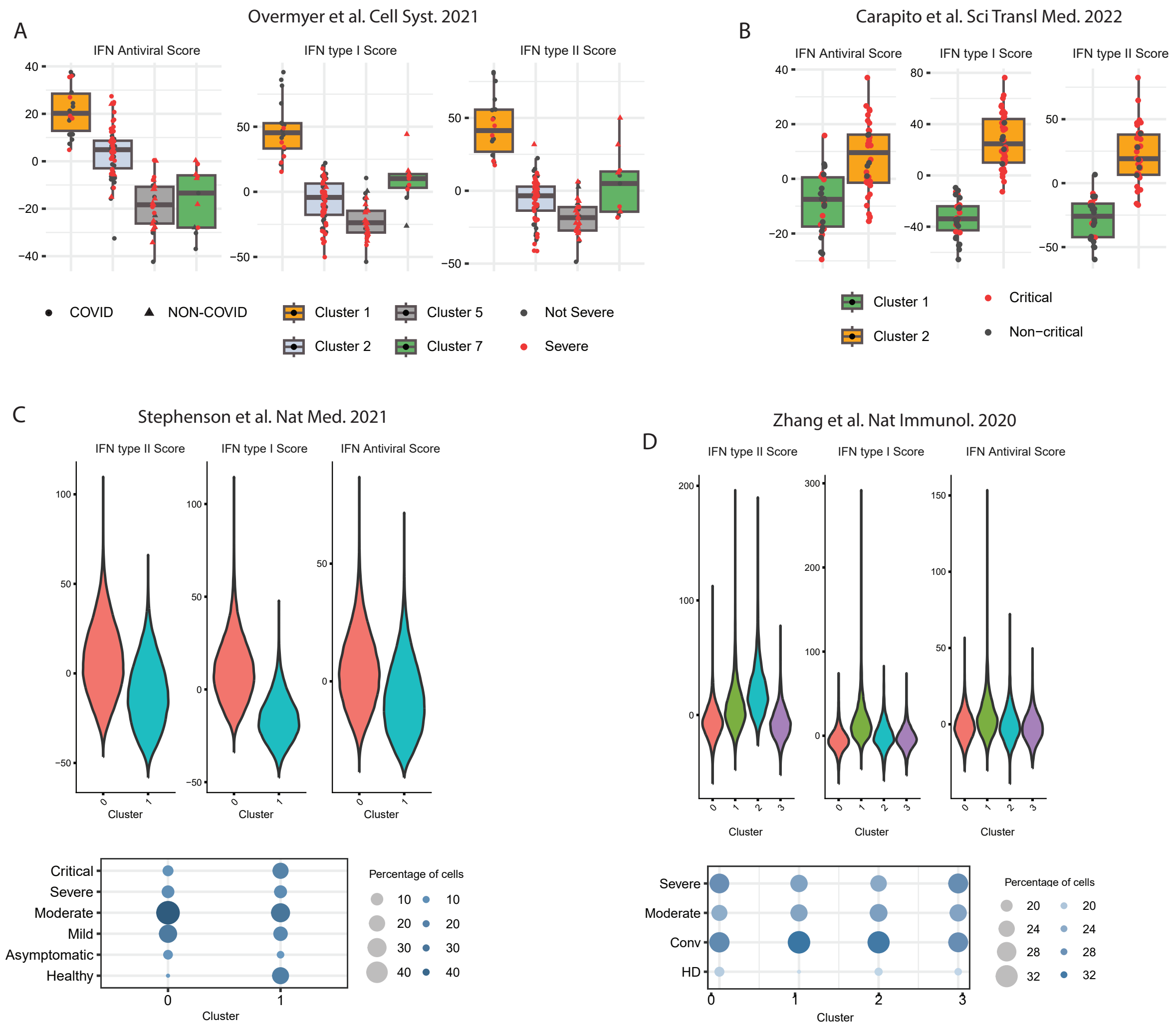


**Supplementary Figure S3:** Independent transcriptomic datasets confirm heterogeneous interferon-stimulated gene (ISG) states across COVID-19 cohorts. External bulk and single-cell transcriptomic datasets were analyzed using the same ISG scoring framework applied in this study to evaluate whether ISG-associated transcriptional heterogeneity could be detected across independent cohorts. (A) Bulk RNA-seq dataset from Overmyer *et al.* [30] showing antiviral, type I interferon (IFN-α/β), and type II interferon (IFN-γ) ISG scores across ISG-defined clusters. Individual samples are displayed as points and colored according to disease severity classification used in the original study (severe vs non-severe). (B) Bulk RNA-seq dataset from Carapito *et al.* [31] analyzed using the same ISG scoring approach. Boxplots display antiviral, type I IFN, and type II IFN ISG scores across the ISG-defined clusters, with individual samples colored by clinical status (critical vs non-critical) as defined by the study. (C) Single-cell RNA-seq dataset from Stephenson *et al.* [32]. Violin plots (top panel) show ISG scores for antiviral, type I IFN, and type II IFN pathways across monocyte clusters based on ISG expression patterns derived from individual cells. (D) Single-cell RNA-seq dataset from Zhang *et al.* [32]. Violin plots (top panel) show antiviral, type I IFN, and type II IFN ISG scores across monocyte clusters based on ISG expression patterns. Bubble plots below panels C and D indicate the proportion of cells originating from different clinical categories, as defined in the respective, contributing to each cluster.

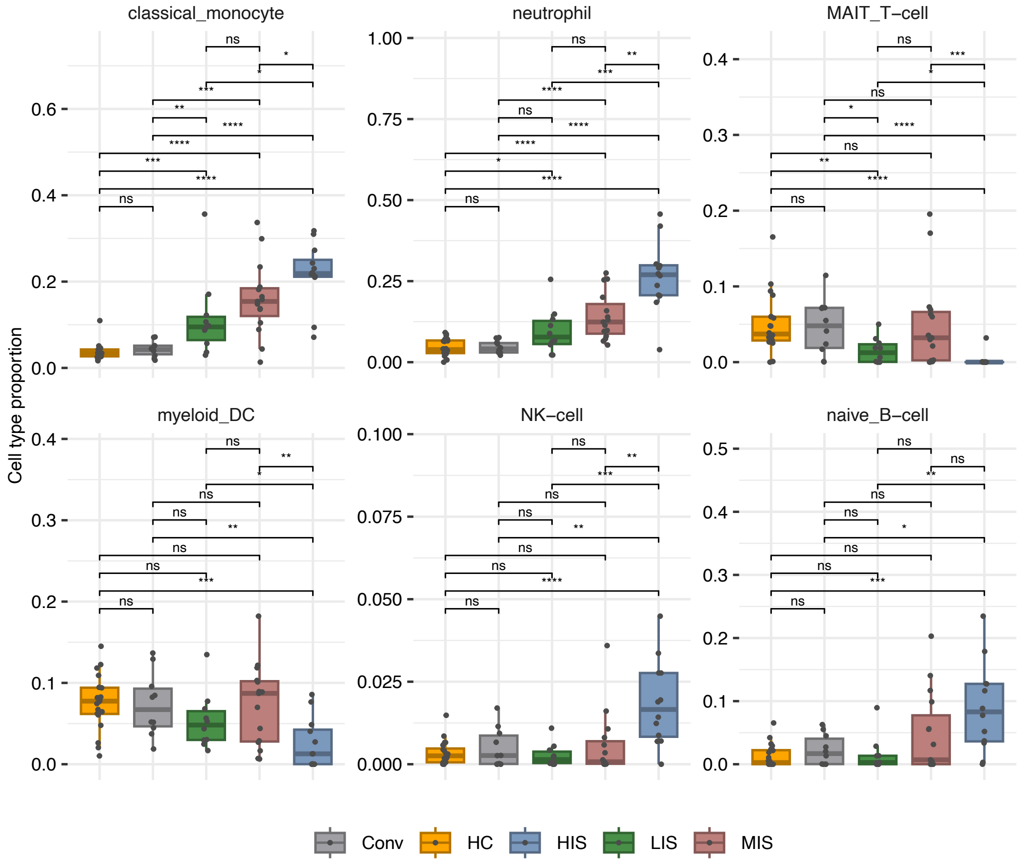


**Supplementary Figure S4:** Box plots showing inferred proportions of selected immune cell types identified by digital cell quantification (DCQ) using EPIC across study groups: healthy controls (HC), convalescent individuals (Conv), low ISG score (LIS), moderate ISG score (MIS), and high ISG score (HIS). Only immune cell populations that showed a statistically significant difference in at least one pairwise comparison after multiple-testing correction (Benjamini–Hochberg adjusted q < 0.05) are included in this figure. Statistical comparisons between groups were performed usi-g the Mann-Whitney U test. For visualization clarity, significance stars displayed in the figure denote unadjusted p-values (p < 0.05, *p < 0.01, **p < 0.001), while the selection of cell types shown was based on adjusted q-values. DCQ-derived cell proportions are shown to provide qualitative insight into systemic immune-state differences rather than absolute immune cell frequencies. Data points are colored based on cluster categorization, with statistical comparisons performed within groups.

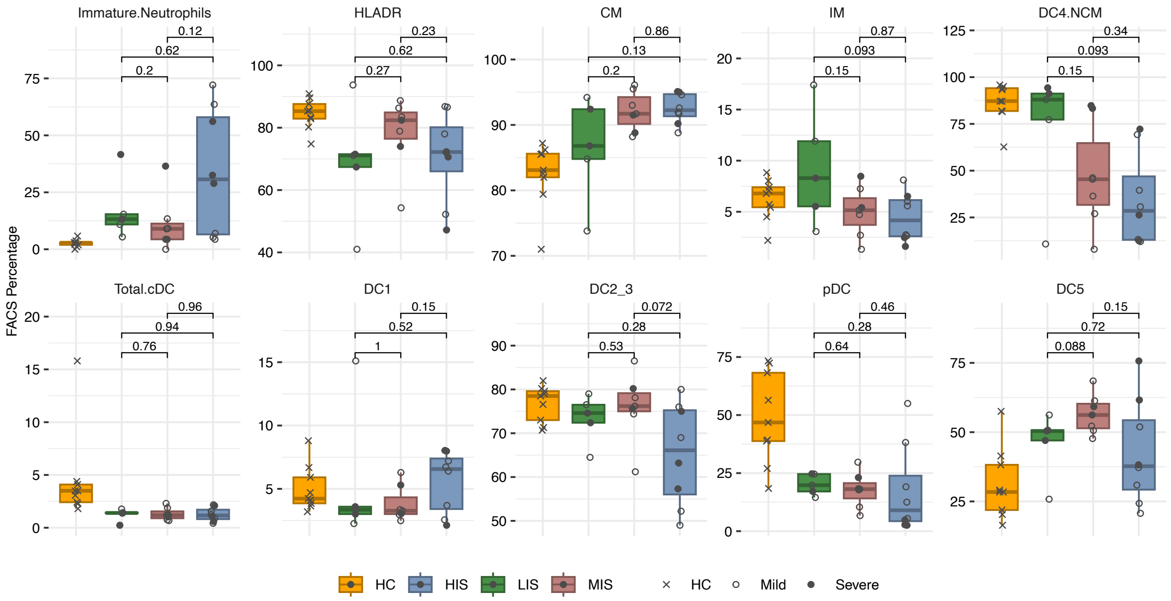


**Supplementary Figure S5: Flow cytometry analysis of myeloid cell populations across ISG-defined clusters.** Box plots representing the frequency of indicated myeloid cell populations identified by flow cytometry in a subset of patients belonging to different ISG-defined clusters **(**HC: n=9, LIS: n=5, MIS: n=7, HIS: n=8**).** The analyzed subsets include immature neutrophils, HLA-DR+ cells, classical monocytes (CM), intermediate monocytes (IM), non-classical monocytes (DC4:NCM), total conventional dendritic cells (cDC), DC1, DC2/3, plasmacytoid dendritic cells (pDC), and DC5. Statistical significance between groups was determined using the Mann-Whitney U test, with p-values indicated above the comparisons.

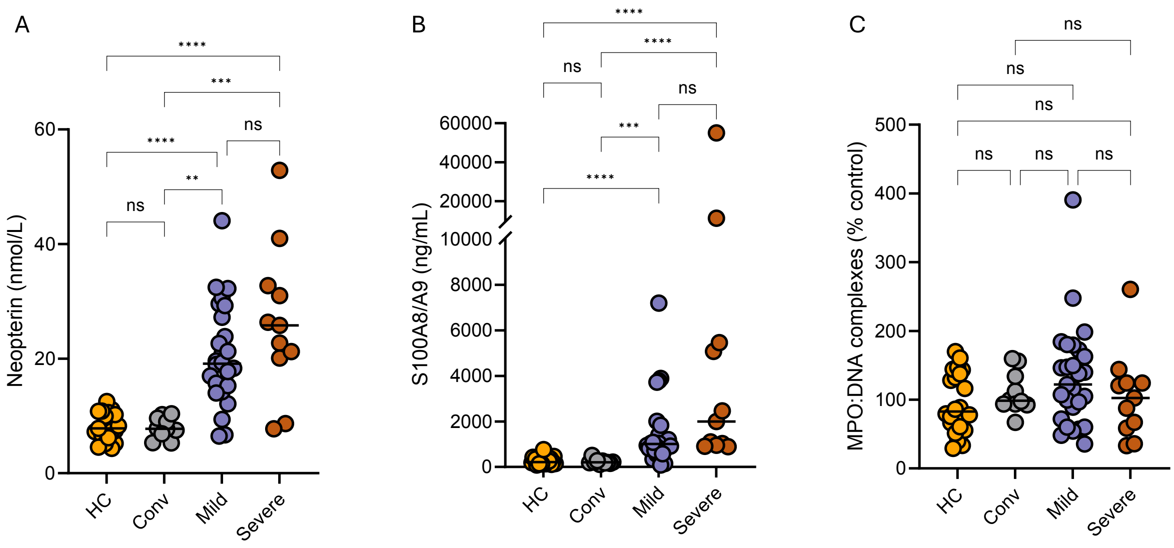


**Supplementary Figure S6: Plasma levels of inflammatory and neutrophil activation markers across disease conditions.** Dotplots displaying plasma levels of (A) Neopterin (nmol/L), (B) S100A8/A9 (ng/mL), and (C) MPO:DNA complexes (% of control) in healthy controls (HC), convalescent individuals (Conv), and COVID-19 patients categorized as mild or severe. Statistical significance was determined using the Mann-Whitney U test, with significance levels indicated as $p<0.05$ (*), $p<0.01$ (**), $p<0.001$ (***), and $p<0.0001$ (****); ns = not significant.

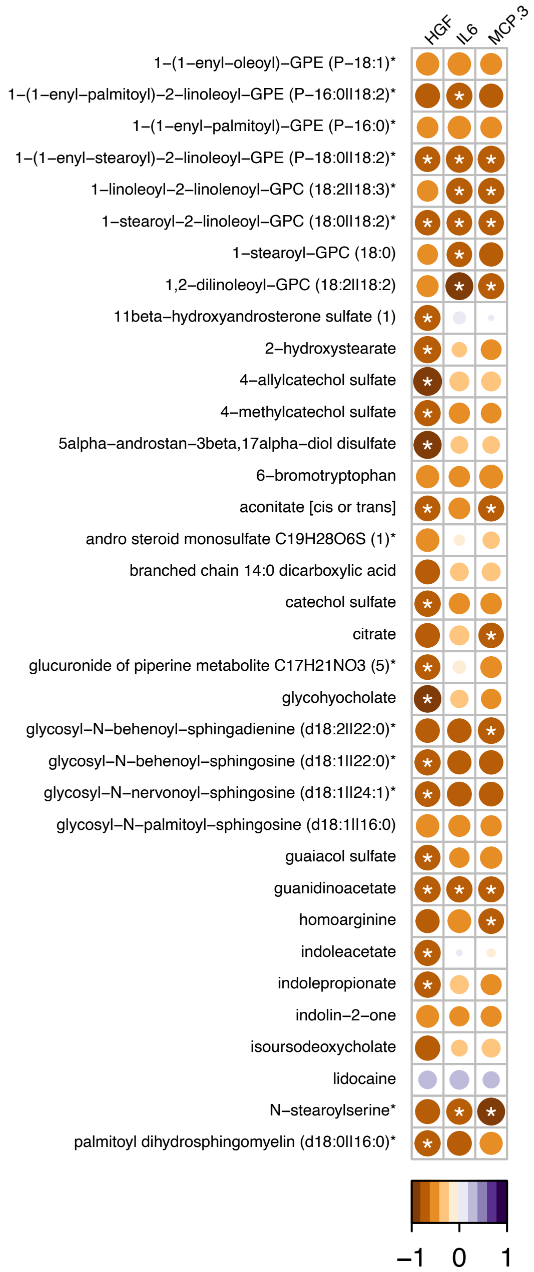


**Supplementary Figure S7: Correlation between metabolite levels and immune activation markers in the HIS cluster.** Heatmap showing Spearman correlation coefficients between significantly altered metabolites and soluble plasma inflammatory mediators (MCP3, HGF and IL6) in HIS patients. The color scale represents correlation strength, with purple indicating positive correlations and brown indicating negative correlations. Asterisks (*) indicate statistical significance (*p* < 0.05).

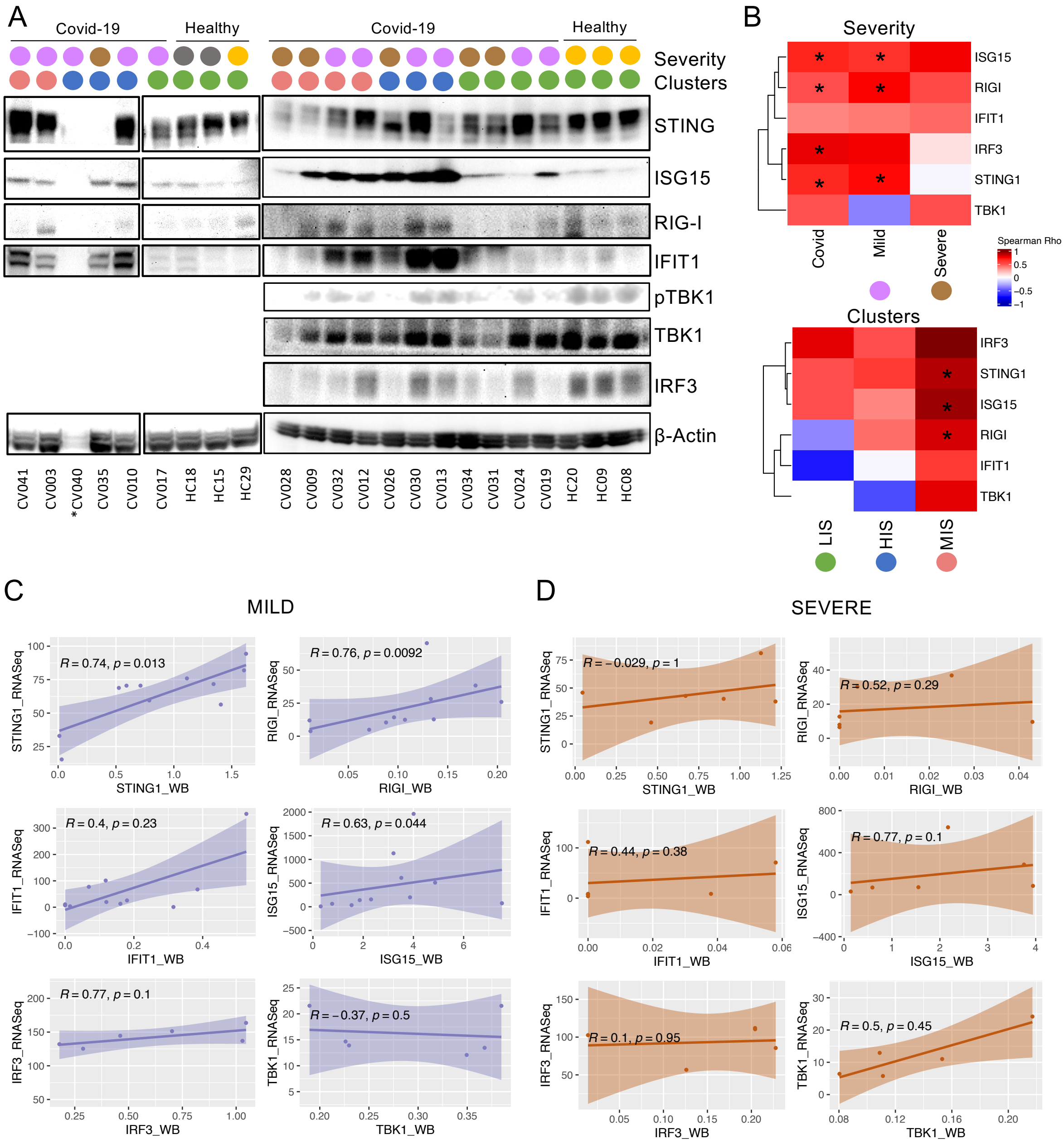


**Supplementary Figure S8:** Western blot for indicated antibodies in PBMCs from Healthy (n=4), Convalescent (n=2), Mild (n=10) and Severe (n=6) patient groups. *CV40 was excluded considered due to low expression of loading control. The severity and the COVID-19 ISG clusters are indicated by their respective colors. pTBK1, TBK1 and IRF3 could not be performed in a subset due to low protein amount **(A)**. Heatmap for correlation between the protein expression levels determined by densitometric quantification of western blot and the mRNA transcript levels determined by the RNAseq in COVID-19 patients categorized based on severity and the ISG-scored clusters. Significant correlations are marked by * **(B)**. Scatterplots for spearman correlation in the mild and the severe group **(C)**.

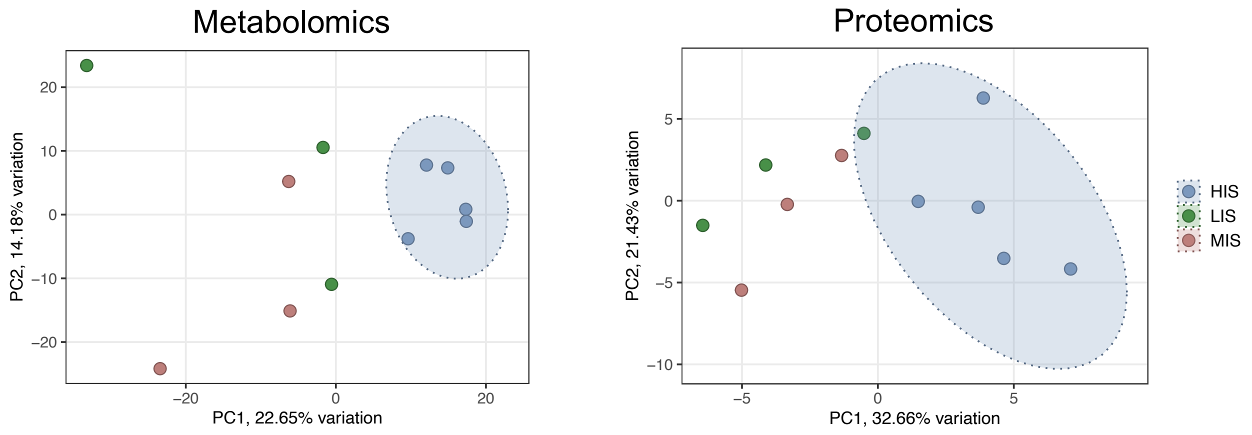


**Supplementary Figure S9:** Distinct systemic molecular states among clinically severe COVID-19 patients across ISG-defined endotypes. Principal component analysis (PCA) of plasma metabolomics (left) and proteomics (right) restricted to patients with severe COVID-19. Samples are colored by ISG-defined endotype: low ISG score (LIS), moderate ISG score (MIS), and high ISG score (HIS). Ellipses represent the 95% confidence interval for each group. Despite comparable clinical severity, severe HIS patients segregated from severe LIS and MIS patients in both metabolomic and proteomic PCA space, indicating distinct systemic molecular profiles. PCA axes are labeled with the percentage of variance explained.

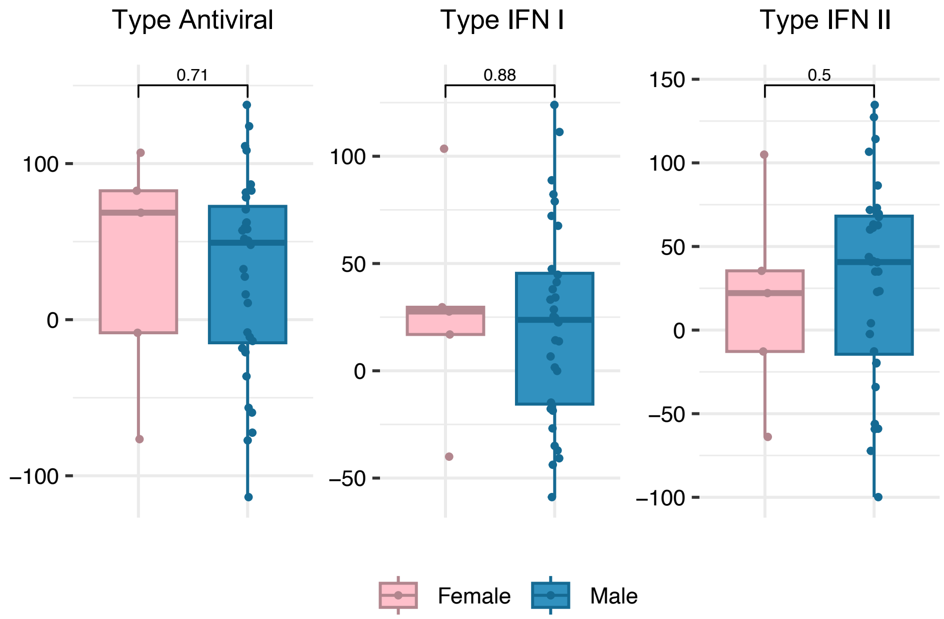


**Supplementary Figure S10:** Sex-associated comparison of systemic interferon-stimulated gene (ISG) expression. Boxplots show antiviral, type I IFN-associated, and type II IFN-associated ISG scores stratified by sex across the COVID-19 cohort. Individual data points represent patients, with boxes indicating median and interquartile range and whiskers denoting 1.5× IQR. Statistical comparisons between female and male patients were performed using the Mann-Whitney U test; exact p-values are indicated above each comparison. No statistically significant differences in ISG scores were observed between sexes across any ISG category, indicating that sex does not substantially influence the magnitude of systemic ISG expression in this cohort.

**Supplementary Tables:**

**Table S1:** List of interferon-stimulated genes (ISGs) used for ISG score calculation.
**Table S2:** Summary of patient-specific ISG scores for type-I, type-II, and antiviral ISGs.
**Table S3:** Plasma levels of IFNs measured using LEGENDplex assay.
**Table S4:** Differentially expressed plasma proteins across ISG-defined clusters.
**Table S5:** List of significantly altered metabolites in ISG-defined clusters.
**Table S6:** Correlation of ISG scores with significantly altered metabolites.
**Table S7:** Digital cell quantification (DCQ) results for immune cell fractions across ISG clusters.
**Table S8:** Plasma levels of S100A8/A9 and Neopterin in ISG-defined clusters.
**Table S9:** Correlation of ISG scores with immune cell proportions.
**Table S10:** Differentially expressed inflammatory mediators (Olink Immuno-oncology panel) in HIS patients with mild vs. severe disease.
**Table S11:** Significantly altered metabolites in HIS patients with mild vs. severe disease.
**Table S12:** Correlation of metabolic alterations with immune activation markers.
